# Transcriptional programs define intratumoral heterogeneity of Ewing sarcoma at single cell resolution

**DOI:** 10.1101/623710

**Authors:** M-M Aynaud, O Mirabeau, N Gruel, S Grossetête, V Boeva, S Durand, D Surdez, O Saulnier, S Zaïdi, S Gribkova, U Kairov, V Raynal, F Tirode, TGP Grünewald, M Bohec, S Baulande, I Janoueix-Lerosey, J-P Vert, E Barillot, O Delattre, A Zinovyev

## Abstract

EWSR1-FLI1, the chimeric oncogene specific for Ewing sarcoma (EwS), induces a cascade of signaling events leading to cell transformation. However, it remains elusive how genetically homogeneous EwS cells can drive heterogeneity of transcriptional programs. Here, we combined independent component analysis of single cell RNA-sequencing data from diverse cell types and model systems with time-resolved mapping of EWSR1-FLI1 binding sites and of open chromatin regions to characterize dynamic cellular processes associated with EWSR1-FLI1 activity. We thus defined an exquisitely specific and direct, super-enhancer-driven EWSR1-FLI1 program. In EwS tumors, cell proliferation was associated with a well-defined range of EWSR1-FLI1 activity; moreover, cells with a high EWSR1-FLI1 activity presented a strong oxidative phosphorylation metabolism. In contrast, a subpopulation of cells from below and above optimal EWSR1-FLI1 activity was characterized by increased hypoxia. Overall, our study reveals sources of intratumoral heterogeneity within Ewing tumors.

## Introduction

Ewing sarcoma (EwS) is a highly aggressive pediatric bone cancer, which is defined by a pathognomonic recurrent somatic mutation - a fusion between the *EWSR1* gene and an *ETS* family member, most frequently the *FLI1* gene (Delattre et al., 1992; Grunewald et al., 2018). This leads to the expression of EWSR1-FLI1, an aberrant and potent chimeric transcription factor. EWSR1-FLI1 can act both as a transcriptional activator and as a repressor, depending on the sequences of DNA binding sites and on the presence of additional co-factors (Bilke et al., 2013; Riggi et al., 2014). EWSR1-FLI1 binds to DNA either at ETS-like consensus sites with a GGAA core motif or at GGAA-microsatellites (GGAA-mSats) which are diverted by EWSR1-FLI1 as *de novo* enhancers (Gangwal et al., 2008; Guillon et al., 2009; Riggi et al., 2014). Through binding to these sites, EWSR1-FLI1 has been reported to act directly or indirectly on many key cellular processes including cell cycle, apoptosis, angiogenesis, metabolism and cell migration (Grunewald et al., 2018; Stoll et al, 2013).

EwS is genetically stable and ranks among tumors with the lowest mutation rates (Brohl et al., 2014; Crompton et al., 2014; Lawrence et al., 2013; Tirode et al., 2014). Indeed, apart from the *EWSR1-FLI1* fusion, EwS harbors only few other recurrent mutations at low frequencies: *TP53* (5-10%), *CDKN2A* (10%) and *STAG2* (15%) (Brohl et al., 2014; Crompton et al., 2014; Grunewald et al., 2018; Huang et al., 2005; Tirode et al., 2014). Despite this remarkable paucity of somatic mutations, EwS is a very aggressive tumor with a strong propensity to progress, metastasize and resist to treatments, suggesting potent adaptation properties of cancer cells. Recent data suggests that epigenetic and transcriptional heterogeneity can play an important role in explaining the mechanisms of such adaptation. Indeed, from a large cohort of EwS tumors, a study on a genome-scale DNA methylation sequencing described consistent DNA hypomethylation at enhancers regulated by EWSR1-FLI1 and strong epigenetic heterogeneity within tumors (Sheffield et al., 2017). Moreover, variable expression of EWSR1-FLI1 was recently suggested as a source of heterogeneity in cell lines and tumors, with a high level of EWSR1-FLI1 expression (EWSR1-FLI1^high^) cells being highly proliferative whereas EWSR1-FLI1^low^ demonstrate a strong propensity to migrate, invade and metastasize (Franzetti et al., 2017). EwS therefore constitutes an appropriate model to investigate how a single somatic driver mutation may impact on critical cell fate decisions ultimately leading to tumorigenesis.

Intratumoral heterogeneity can now be investigated at the single cell level through single cell ‘omics’ technologies that enable to explore in great details the cell-to-cell variations in gene expression (Baslan and Hicks, 2017). These approaches can help characterize the origins of genetic and non-genetic heterogeneity which can modulate cell response to exogenous and endogenous factors such as the activation of cancer driver genes (Almendro et al., 2013). Such approaches can also decipher essential bi-or multi-modalities in the distribution of expression of the genes regulating the cell fates (Shalek et al., 2013) or the interplay between progression through the cell cycle and the action of signaling and/or differentiation pathways, which can hardly be addressed through bulk-cell analysis (Buettner et al., 2015).

Here, we first used single cell analysis to explore the dynamics of EWSR1-FLI1-related expression changes at the single cell level using a time course upon the EWSR1-FLI1 induction. Ewing cell transcriptomic profiles were also compared with a set of single cell profiles from other reference systems chosen by various aspects of similarity to the Ewing cell system: being either time series experiments, cells corresponding to EwS cell-of-origin or cells of various tumor types. This analysis was combined with the exploration of the epigenetic changes. Overall, this approach allowed us to distinguish generic dynamics of transcriptional changes that are shared by most scrutinized systems from system-specific, and particularly EwS-specific, dynamics. These components were then used to investigate single cells from Ewing tumors. This two-steps approach illuminates the heterogeneity of Ewing tumors, distinguishing different cell populations based on EWSR1-FLI1 activity, proliferation and metabolic characteristics.

## Results

### Experimental design for collecting EwS single cell RNA-sequencing profiles

In order to explore the dynamics of individual cell transcriptomes under *EWSR1-FLI1* expression, we used the previously developed A673/TR/shEF inducible cellular model derived from the A673 EwS cell line where the expression of *EWSR1-FLI1* can be modulated through a doxycycline-controlled shRNA (Carrillo et al., 2007). Following down-modulation of *EWSR1-FLI1* (*EWSR1-FLI1*^*l*ow^) by 7 days of doxycycline (DOX) treatment, we performed a time course experiment after removal of DOX from the medium leading to *EWSR1-FLI1* re-expression. We made single-cell transcriptome measurements at 7 time points (days 7: *EWSR1-FLI1*^*l*ow^, 9, 10, 11, 14, 17 and 22: *EWSR1-FLI1*^*high*^) (Figure 1A). We also tested *in vivo* the impact of *EWSR1-FLI1* on gene expression. From A673/TR/shEF xenografts in SCID mouse, single-cell RNA-sequencing (scRNA-seq) was conducted without DOX (*EWSR1-FLI1*^high^) and after 7 days of DOX treatment (*EWSR1-FLI1*^*l*ow^). The modulation of EWSR1-FLI1 protein expression was confirmed by western blot (Figure 1B). In addition, the expression of CD99 at the membrane and the nuclear expression of LOX, surrogate markers of *EWSR1-FLI1* activation and repression respectively, was confirmed by immunohistochemistry (IHC) (Figure 1B). We also conducted scRNA-seq experiments on 3 Ewing patient derived xenografts (PDX), established in the laboratory by implantation of tumor samples in SCID mice. PDX-83 and PDX-84 expressed *EWSR1-FLI1* type I fusion and PDX-111 harbors *EWSR1-FLI1* type X fusion. Finally, we profiled two primary cultures of mesenchymal stem cells (MSCs), the likely EwS cell-of-origin (Tirode et al., 2007).

**Figure 1.**
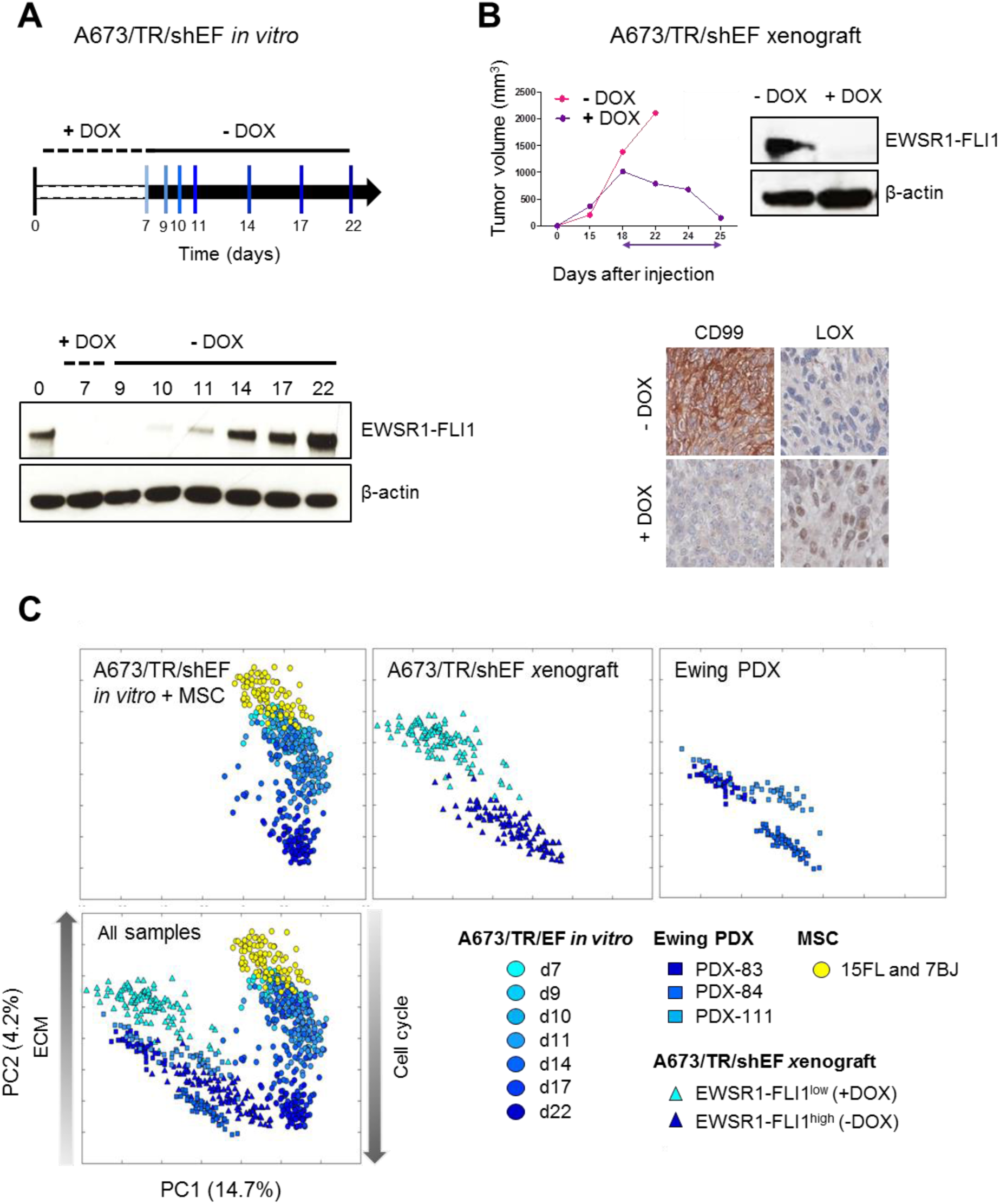
Experimental design for collecting EwS scRNA-seq profiles. **A.** A673/TR/shEF *in vitro* - Scheme of the time-resolved induction of DOX-regulated *EWSR1-FLI1* and western blot. After 7 days of DOX treatment, cells are extensively rinsed, grown in a DOX-free medium then collected at 2 days (d9), 3 days (d10), 4 days (d11), 7 days (d14), 10 days (d17) and 15 days (d22) and isolated using the Fluidigm™ C1 system for scRNA-seq. **B.** A673/TR/shEF xenograft. A673/TR/shEF were injected sub-cutaneously in SCID mice. After 18 days, mice were separated in two groups, one with DOX added to the drinking water for 7 days and the other group without DOX. EWSR1-FLI1 expression was controlled by western blot. IHC with positive (CD99) and negative (LOX) EWSR1-FLI1 surrogate markers are shown. **C.** PCA of single cells EwS datasets. Cells are indicated by colored circles, from d7 (light blue circles) to d22 (dark blue dot), A673/TR/shEF xenograft cells at d0 (dark blue triangle) and d7 (light blue triangle), 3 Ewing PDXs (blue square), MSCs (yellow circles).

Unsupervised analysis of the Ewing single-cell transcriptomic data by Principal Component Analysis (PCA), clearly separated *in vitro* (A673/TR/shEF time series & Ewing MSCs) (Figure 1C, upper left panel) and *in vivo* datasets (A673/TR/shEF xenograft & Ewing PDXs) (Figure 1C, middle and right upper panel) along the first principal component (PC1, 14.7% of explained variance) (Figure 1C). The second principal component (PC2, 4.2% of explained variance) projection revealed the effect of *EWSR1-FLI1* activation on transcriptomic dynamics. For A673/TR/shEF time series, *EWSR1-FLI1*^*l*ow^ cells (d7) were grouped close to MSCs and clearly separated from *EWSR1-FLI1*^high^ cells (d22). As early as the second (d9) and the third (d10) days of *EWSR1-FLI1* re-expression, the distribution of single cell transcriptomes is already significantly different from *EWSR1-FLI1*^*l*ow^ cells (d7). 4 and 7 days after re-induction (d11 and d14) represent intermediate (between *EWSR1-FLI1*^low^ and *EWSR1-FLI1*^high^) states of single cell transcriptome distributions. Finally, at d17 most of the single cells converge to the *EWSR1-FLI1*^high^ state of d22. Similarly, *EWSR1-FLI1*^low^ and *EWSR1-FLI1*^high^ states of A673/TR/shEF xenografts could be clearly distinguished. All *EWSR1-FLI1*^high^ cells, including the 3 PDXs, converge together in the transcriptomic space (Figure 1C, All samples). The first component was not significantly enriched with any gene ontology gene set, while the second principal component was associated by functional enrichment analysis to EWSR1-FLI1 modulated genes (taken from Kinsey et al., 2006), positive correlation, *p* < 10^−65^), Cell Cycle (GO:0007049, positive correlation, *p* < 10^−26^), and Extracellular Matrix (ECM) Organization (GO:0030198, negative correlation, *p* < 10^−25^).

We also checked the single cell expression dynamics of 8 genes known to be directly modulated by *EWSR1-FLI1* (up-regulated genes: *PRKCB, LIPI, CCND1, NR0B1*; down-regulated genes: *IGFBP3, IL8, LOX, VIM*) (Figure S1). These results confirmed consistent re-induction dynamics of *EWSR1-FLI1*. Single cell expression of these genes highlights early and late responsive cells to *EWSR1-FLI1* re-expression at any given time point (Figure S1) (Hancock and Lessnick, 2008).

Collectively, these analyses show that these Ewing single cell transcriptome datasets recapitulated the main results found previously in bulk expression measurements in similar biological systems. However, just as in the bulk data, this standard multivariate analysis did not allow to distinguish processes directly related to EWSR1-FLI1 specific transcriptional activity from generic biological processes (cell cycle, ECM organization, etc…), indirectly modified following *EWSR1-FLI1* expression.

### Joint deconvolution of multiple scRNA-seq datasets into independent components

In order to create a negative control to the Ewing-specific datasets and evaluate the specificity of the sources of cellular heterogeneity, we jointly normalized and merged the Ewing-specific single cell datasets together with several other single cell datasets generated in-house or obtained from public resources (Patel et al., 2014; Trapnell et al., 2014). Altogether we analyzed 1,964 single cell transcriptomic profiles from 8 different datasets (Table S1). A t-distributed stochastic neighbor embedding (t-SNE) plot of all cells is shown in Figure S2. The different Ewing-specific datasets are grouped together, separated from the other datasets. In both *in vitro* and xenograft cases, cells in which the *EWSR1-FLI1* oncogene has been induced converge to the same area at the center of the plot. Cells of mesenchymal origin (MSC and myoblasts) are localized close to each other in the plot.

We applied Independent Component Analysis (ICA) in order to decompose the heterogeneity of scRNA-seq profiles into a relatively small number (30, as it was close to the estimated optimal number, see Methods) of independently acting factors or independent components (ICs). The rationale for choosing this approach is that ICA can in principle deconvolute signals correlated to a common hidden factor, since the covariance matrix was made unity before application of ICA by data whitening (Zinovyev et al., 2013). For each IC, the analysis associated a weight for each gene (collectively denoted as metagene) and a score for each sample (denoted as metasample) (Tables S2 and 3).

Correlating the computed 30 meta-samples to the binary vectors describing different cell subsets enabled us to distinguish generic and cell type-specific independent sources of heterogeneity (Figure 2A and Table S2) including Ewing specific independent components (IC10, IC30) as well as components specific to other cell types (Figure 2A). In addition, ICA deconvolution leads to the identification of components not specific to any cell type, whose correlations with the cell subsets binary vectors were small (those are located in the right bottom part of the Figure 2A).

**Figure 2.**
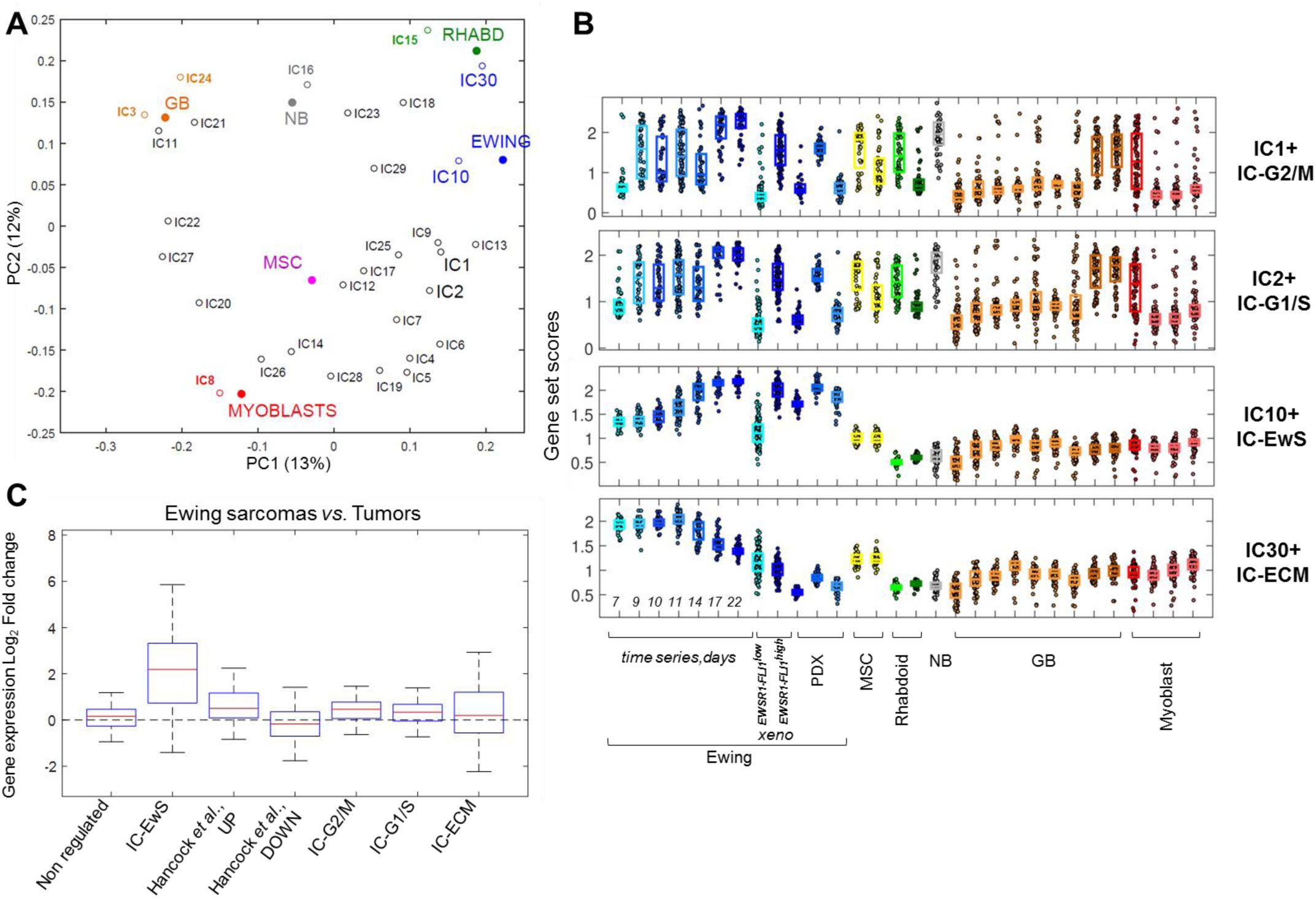
Deconvoluting the cell cycle phases and specific transcriptional program of EWSR1-FLI1 activity. **A.** PCA plot visualizing the matrix of correlations computed between independent components (ICs) and the binary vectors distinguishing different groups of cells: Ewing (blue), Neuroblastoma, NB (grey), Rhabdoid, RHABD (green), Myoblast (red), Glioblastoma, GB (orange). **B.** Gene set score distribution across all cell datasets for four selected ICs (IC1+: IC-G2/M, IC2+: IC-G1/S, IC10+: IC-EwS and IC30+: IC-ECM). The scores are computed as average value of the gene expression of the most variable genes in the set (see Materials and Methods). For Ewing dataset, blue scale illustrate EWSR1-FLI1 level of expression: from EWSR1-FLI1^low^ (light blue) to EWSR1-FLI1^high^ (dark blue). For Rhabdoid dataset, green scale shows SMARCB1^−^ (light green) and SMARCB1^+^ (dark green). For GB dataset, GB cell line are in light orange and GB tumors are in dark orange. For the myoblast dataset, red color scale illustrate the myoblast differentiation time course. **C.** Specificity of IC-EwS gene set for EwS. Gene expression analysis is applied on a cohort of 22,956 non EwS tumors and 156 EwSs (all measured by Affymetrix HG-U133Plus 2.0 array). Box plot of gene expression log_2_ fold change of EwS *vs*. other tumors of the non regulated genes (n = 100), IC-EwS genes (n = 220), the up-(n = 503) and down-regulated genes (n = 293) described by Hancock and Lessnick, 2008(Hancock and Lessnick, 2008), the IC-G2/M (n = 212), IC-G1/S (n = 291) and IC-ECM (n = 252) genes.

### Generic and *EWSR1-FLI1* specific components

We then looked for biological processes that could be associated with each of the ICs. For this we defined two sets of top contributing genes for each component, one with positive weights and one with negative weights (denoted as ICx+ and ICx-respectively where x denotes the component number and +/- indicates the long and short tail of the weight distribution, respectively), using a threshold of 5 standard deviations roughly corresponding to a statistical significance of 1%. On these, we then performed gene set enrichment analyses using the Toppgene suite (Chen et al., 2009) (online table: http://bioinfo-out.curie.fr/projects/sitcon/mosaic/toppgene_analysis/). This analysis highlighted associations with various generic biological processes, some having remarkably strong enrichment scores, and led us to focus on 4 gene sets: IC1+, IC2+, IC10+ and IC30+. Thus, IC1+ was associated with chromosome segregation (GO:0007059, *p* = 10^−80^) and mitotic nuclear division (GO:0007067, *p* = 10^−80^), IC2+, with DNA replication (GO:0006260, *p* = 10^−69^) and IC30+, with extracellular matrix organization (GO:30198, *p* = 10^−16^, 10^−25^, and 10^−14^, respectively). IC10+ was not strongly associated with any biological process. We matched the IC1+ and IC2+ scores to two recently established transcriptomic signatures for the specific phases of the cell cycle (Giotti et al., 2017), and found strong match between IC1+ and G2/M score and between IC2+ and G1/S score. Accordingly to this analysis, we will refer to IC1+, IC2+, IC30+ gene sets as IC-G2/M, IC-G1/S and IC-ECM correspondingly, for the sake of clarity (Figure S3).

IC10+ gene set was interpreted as highly Ewing sarcoma-specific; therefore, further we refer to it as IC-EwS. The IC-EwS list (220 genes) was highly enriched in “Genes up-regulated in mesenchymal stem cells (MSC) engineered to express EWSR1-FLI1 fusion protein” (Riggi et al., 2008) (*p* = 10^−102^) and several other Ewing-related transcriptomic signatures from MSigDB C2 collection (targets of EWSR1 ETS fusions up (Miyagawa et al., 2008), *p* = 10^−50^; targets of EWSR1-FLI1 fusion (Hu-Lieskovan et al., 2005), *p* = 10^−48^; Ewing family tumor (Staege et al., 2004), *p* = 10^−25^) and to a lesser extent with the “Ewing sarcoma disease” gene set (C3536893 entry in DisGeNET database (Pinero et al., 2015), *p* = 10^−9^). Unlike previously reported EwS-related gene signatures, IC-EwS was not enriched in cell cycle-related reference gene sets.

We then assigned a gene set activation score to each cell regarding the different ICs (an average expression of most variable genes in the gene set, see Methods). The score distributions allowed to make the following conclusions: (1) IC-G2/M and IC-G1/S scores are distributed across all datasets peaking in the states that can be associated to active proliferation (Figure 2B); (2) within each dataset IC-G2/M and IC-G1/S scores are highly variable; (3) IC-EwS and IC-ECM high score values are almost exclusively associated to Ewing cells; (4) IC-EwS, IC-ECM scores clearly distinguish EWSR1-FLI1^high^ and EWSR1-FLI1^low^ cell states and change monotonically with time, increasing or decreasing, respectively. This is observed in the *in vitro* and xenograft inducible cellular systems. Figure 2B visualizes IC-G2/M, IC-G1/S, IC-EwS, IC-ECM scores across the studied datasets.

To further test the specificity of IC-EwS and IC-ECM gene expression in EwS, we performed gene expression analysis in a combined cohort of 23,112 samples including 156 EwSs and 22,956 other tumors (Gene investigator(Hruz et al., 2008)). The IC-EwS gene set strikingly discriminated EwS from all other samples (Figure 2C), better than the gene signature previously defined by transcriptomic data meta-analysis and containing genes regulated by *EWSR1-FLI1* and enriched in EwSs(Hancock and Lessnick, 2008). This analysis also showed that the IC-ECM gene set is not specific to EwS (Figure 2C).

Altogether, these data highlight that the IC-EwS gene set is highly specific to EwS both in model systems (cell line and PDX) and in tumors.

### Characterization of the specific EWSR1-FLI1 activity signature

To further characterize the IC-EwS signatures, we performed EWSR1-FLI1 ChIP-seq on A673/TR/shEF at d7, d9, d10, d11, d14 and d17, following the experimental design used to generate the A673/TR/shEF time series (Figure 1A). EWSR1-FLI1-specific peaks (EF-peaks) were defined as peaks that significantly varied upon EWSR1-FLI1 re-expression (*p* < 0.005).

For each gene we calculated the distance between the transcription start site (TSS) to the nearest EF-peak. We then compared the distribution of these distances for genes of the various ICs to the distribution of distances of a set of 1,000 control genes that are not regulated by EWSR1-FLI1. As shown in Figure 3A, this distance is significantly smaller for IC-EwS genes (*p* = 10^−29^) as compared to other ICs and to the reference Hancock *et al*. EWSR1-FLI1 signature. Indeed, we observed a highly significant enrichment in percentage of genes with EF-peaks between 0-100 kb from TSS for IC-EwS (38% of genes) as compared to “non-regulated” genes (<10%), from which we concluded that many of IC-EwS genes are likely to be directly regulated by EWSR1-FLI1 (Figure 3B). A slight enrichment was also observed for the 100-200 and 200-300 kb ranges (12%) but not for distances > 300 kb from TSS. TSS of the Hancock *et al*. and of the IC-ECM genes are also slightly significantly closer to EF peaks than controls but with a much less significant statistical association (Figure 3A).

**Figure 3.**
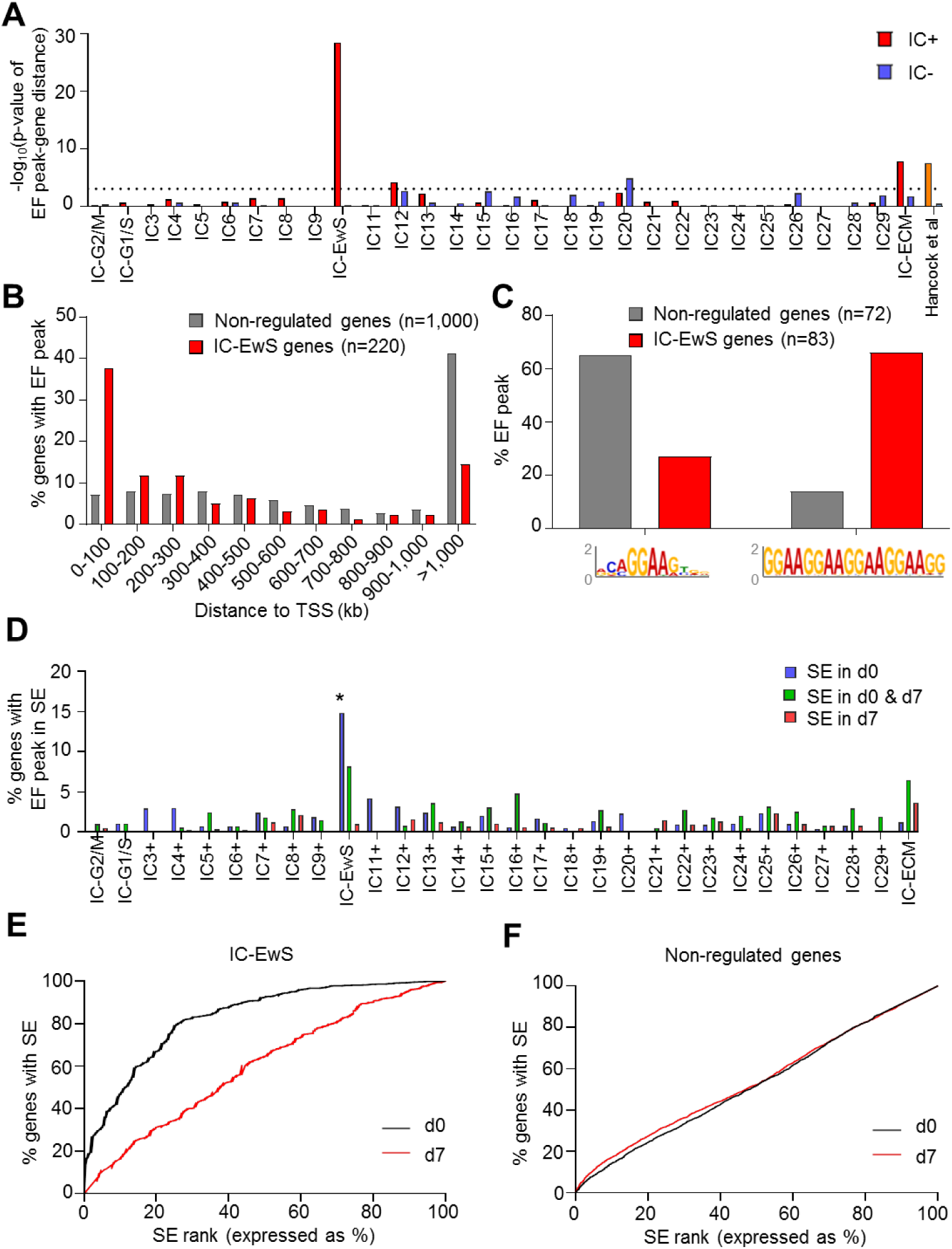
Characterizing the IC-EwS gene set. **A.** Barplot showing for each IC the Log(1/*p*-value) of comparisons of the “gene to EF peak” distance as compared to control genes (“Non-regulated genes” (n = 1,000)). IC+ are in red and IC-in blue. The Hancock *et al EWSR1-FLI1* up and down signatures(Hancock and Lessnick, 2008) are also shown. The dotted line indicates the significant threshold for the *p*-value according to Bonferroni correction (*p* = 8.10^−4^). **B**. Comparison of the enrichment of genes with EF peaks in IC-EwS (n = 220) depending upon the distance to TSS as compared to control genes (n = 1,000). **C**. Analysis of the two types of EF peaks (ETS and GGAA-mSats≥4) in control and IC-EwS genes. Only EF peaks <100 kb of TSS are considered, *i*.*e*. 72 peaks for control and 83 for IC-EwS genes. **D**. Percentage of genes with EF peaks in super-enhancers (as defined by the ROSE software) for the different ICs. The star indicates the group with is significantly different from the control (Fisher test with Bonferroni correction, *p* = 5.10^−4^). **E**. Super-Enhancer rank curve showing enrichment of top super-enhancers (low rank) in IC-EwS genes at d0 (black line) as compared to d7 (red line). **F**. Super-Enhancer rank curve for the control geneset.

Direct EWSR1-FLI1 target binding sites are shown to be either *bona fide* ETS motifs or GGAA-mSats (Gangwal et al., 2008; Guillon et al., 2009; Riggi et al., 2014). We used FIMO (Grant et al., 2011) to analyze the occurrences of these two motifs in EF-peaks located less than 100 kb from TSS of IC-EwS genes (n = 83/220) as compared to control genes (n = 72/1000). While most EF-peaks related to control genes were ETS sites, most EF-peaks of IC-EwS genes contained GGAA-mSats with at least 4 repeats (GGAA-mSats≥4) (Figure 3C).

We also performed ChIP-seq analysis of H3K27ac histone mark to map active chromatin regions at d0 (EWSR1-FLI1^high^) and d7 (EWSR1-FLI1^low^). We observed that 91% of EF-peaks are associated with H3K27ac marks at d0, in agreement with EWSR1-FLI1 being a pioneer factor for chromatin remodeling (Boulay et al., 2017). When considering only EF-peaks localized in super-enhancers (SEs) regions, as defined by the ROSE algorithm (Loven et al., 2013; Whyte et al., 2013), we can define SE associated with EF-peak at d0, at d7 and at both time points. We observed that SEs defined at d0 and containing EF-peaks are enriched in the IC-EwS set of genes (*p* < 10^−5^) (Figure 3D). Moreover, it appeared that the IC-EwS associated SEs ranked among the strongest SEs (Figure 3E). Such an association is specific for IC-EwS as it was observed neither for control genes (Figure 3F) nor for any other ICs (data not shown).

Altogether, these analyses allowed us to define the IC-EwS signature as strongly enriched in EWSR1-FLI1 direct target genes. These genes are associated to EF-peaks that (1) significantly vary upon EWSR1-FLI1 expression, (2) are significantly closer to the TSS, (3) are enriched in GGAAmSat ≥4, (4) are significantly enriched in potent super-enhancers. Given all the analysis steps described above, we defined a set of 83 genes that have most of these characteristics and hence appear to be excellent candidates for being EWSR1-FLI1 direct targets, playing key roles in EwS oncogenesis (Figure 4).

**Figure 4.**
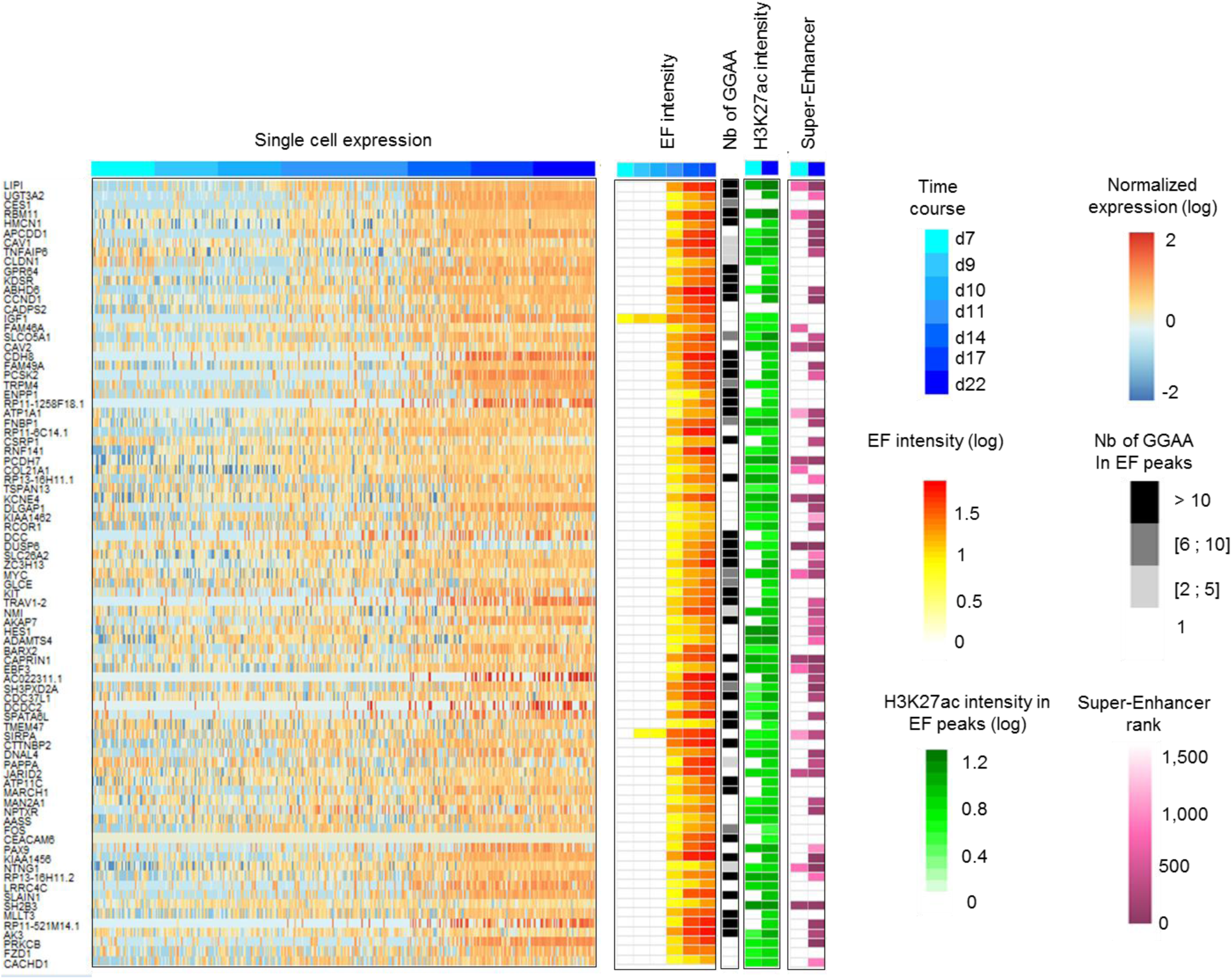
The 83 genes identified as likely direct EWSR1-FLI1 targets. From left to right: heatmap of their expression in single cell normalized expression center scaled and winsorized (5%) in A673/TR/shEF time course from EWSR1-FLI1^low^ (d7, light blue) to EWSR1-FLI1^high^ (d22, dark blue), heatmap of EF-peak intensities in A673/TR/shEF time course from EWSR1-FLI1^low^ (d7, white) to EWSR1-FLI1^high^ (d17, red), number of GGAA in EF-peaks (grey scale), heatmap of H3K27ac histone mark co-localized with EF peaks (from light green to dark green) and of super-enhancer ranking in A673/TR/shEF d0 and d7 (from dark purple to light purple). The 83 genes are ranked by their weight in IC-EwS.

### Unraveling heterogeneity of Ewing tumors at single cell level in tumors

We then investigated whether the afore-mentioned signatures, mostly defined by *in vitro* systems, may be informative to explore the structure of large single cell datasets obtained from *in vivo* samples. 5 new patient-derived xenografts (PDX-352, PDX-861, PDX-856, PDX-184 and PDX-1058), from *EWSR1-FLI1*-positive EwSs were profiled using the 10x genomics sequencing platform. After quality checks and removal of profiles corresponding to dead cells, a total of 3,595, 1,245, 604, 1,245 and 1,742 scRNA-seq profiles was obtained for PDX-352, PDX-861, PDX-856, PDX-184 and PDX-1058, respectively. In order to visualize distances between individual cell transcriptomes, we used the SPRING web-based data visualization interface based on the application of a force-directed graph layout to the graph of similarity between full transcriptomic profiles of individual cells (kNN graph) (Weinreb et al., 2018). When IC-EwS score, which can be considered as a direct assessment of EWSR1-FLI1 transcriptional activity, was mapped onto the SPRING layout in all PDXs, we observed that this signature largely contributes to the intratumoral heterogeneity (Figure 5A and Figure S4A-D). In all PDXs, IC-G2/M and IC-G1/S define specific groups of cells that form a loop-like structure, most probably reflecting the transcriptional dynamics of the cell cycle program (Figure 5A-B, Figure S4A-D).

**Figure 5.**
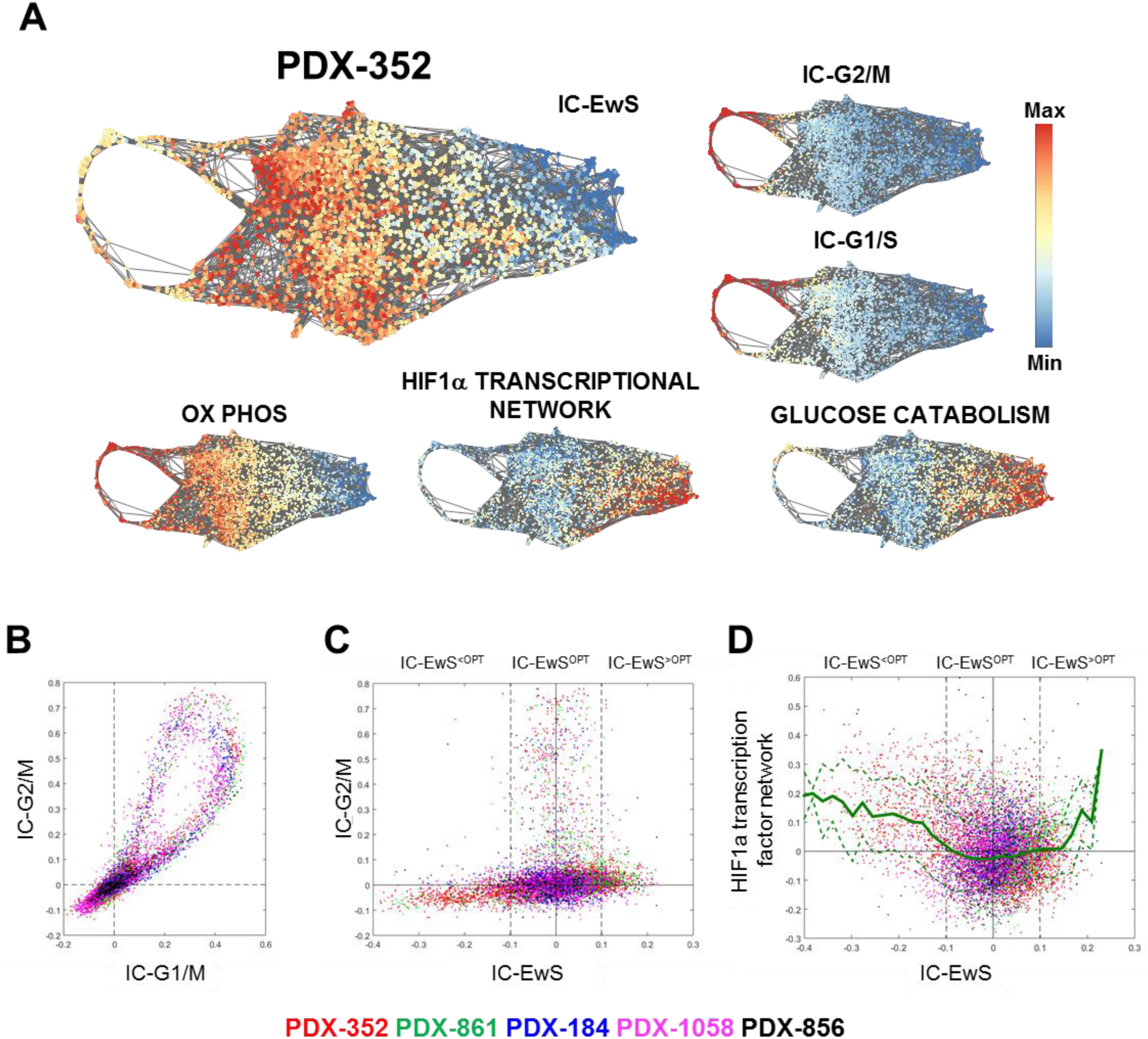
Quantification of PDXs heterogeneity based on identified transcriptional signatures. **A.** SPRING representation of the kNN graphs (k = 5) for PDX-352 dataset. The scores are either gene set (top contributing genes) scores of the ICs computed for model systems (IC-EwS recapitulating the transcriptional program of EWSR1-FLI1, IC-G2/M and IC-G1/S recapitulating the transcriptional programs of the cell cycle) or the scores of the reference gene sets recapitulating a biological function contributing to the intratumoral heterogeneity. An interactive version of this Fig. is available at https://goo.gl/XqCKuC **B.** Visualization of the cell cycle transcriptional program, used to define the proliferating cells of the 5 EwS PDXs. The scores were centered to have zero median for non-proliferating cells, separately for each PDX. **C.** Plot visualizing connection between IC-EwS and IC-G2/M scores shows that proliferative cells tend to appear in an optimal range IC-EwS^OPT^ of IC-EwS values. IC-EwS scores were centered to have zero median for proliferating cells, in each PDX. **D.** Plot showing connection between IC-EwS and HIF1α transcription factor network score. The HIF1α scores were centered to have zero median, in each PDX. In panels **B**,**C**,**D**, solid line shows local average of the score, and the dashed lines indicate one standard deviation interval.

An attempt was made to merge together the 5 PDXs transcriptomic datasets, but their transcriptomes remain too specific to be directly compared even when adjusted for the variations in library size. Instead, we found that the score distributions for specific set of genes in our focus of study can be easily aligned (Figure 5A). Thus, using joint analysis of 5 PDXs, we found that it is plausible to suggest existence of an optimal range of intermediate IC-EwS values to activate cell proliferation (Figure 5C). We define such a range as a window of IC-EwS scores containing 95% of proliferative cells, identified accordingly to IC-G1/S or IC-G2/M scores (Figure 5C). Below and above this range we observe only a small number of cells in proliferative state. By using Kolmogorov-Smirnov test, we verified that this observation can not be a random sampling effect caused by the globally higher density of cells in this defined range (*p* < 10^−22^).

We further investigated which biological factors can potentially explain the lack of proliferation outside the optimal window of EWSR1-FLI1 activity estimated through the IC-EwS score in PDXs. For each PDX separately, we defined IC-EwS^OPT^ as the group of cells whose IC-EwS scores fall into the optimal proliferation range, an interval containing a majority of IC-EwS scores from proliferative cells. Other cells, characterized by IC-EwS scores below and above the optimal range, were respectively labeled as IC-EwS^<OPT^ and IC-EwS^>OPT^. Cells of both of these types were relatively rare (in average, for 5 PDXs, IC-EwS^<OPT^ and IC-EwS^>OPT^ groups were 10% and 8% of the total cell number, correspondingly).

First, we performed GSEA analysis, comparing IC-EwS^OPT^ cells with the IC-EwS^<OPT^ groups. We found that the strongest up-regulated signal characterizing the cells outside the optimal window was related to hypoxia (HALLMARK_HYPOXIA and PID138045 “HIF1α transcription factor network” gene sets from MSigDB, NES = 5.2, 3.16, *p* < 10^−3^) and glucose catabolism (GO:0006007 “glucose catabolic process”, NES = 3.74, *p* < 10^−3^). By contrast, cells residing inside the optimal range for proliferation were characterized by a high oxidative phosphorylation (HALLMARK_OXIDATIVE_PHOSPHORYLATION gene set from MSigDB, NES = 4.3, *p* < 10^−3^). In all PDXs, we observed that most of IC-EwS^<OPT^ cells were characterized by increased glucose catabolism score (for example, Figure 5A). Inspection of hypoxia score in all PDXs showed that it highlights a compact subgroup of IC-EwS^<OPT^ cells (see characteristic example in Figure 5A and D, and all other PDXs in Figure S4A-D). In all PDXs, this subgroup of IC-EwS^<OPT^ cells highly expressed hypoxia-related markers, such as *ALDOA, CA9, NDRG1, VEGFA, ADM, BNIP3* and *NRN1* (all members of the “HIF1α transcription factor network” pathway) (Figure S5).

A relatively rare population of IC-EwS^>OPT^ cells was also characterized by a consistent increase of many genes from the HIF1α transcription factor network, especially towards the maximally observed IC-EwS scores (see Figure 5D). This signal was, however, not the strongest one among others, being masked and dominated by the increased expression of IC-EwS signature genes, in IC-EwS^>OPT^ (which is expected from the definition of IC-EwS^>OPT^). When the IC-EwS score was regressed out from the expression of all genes, only the hypoxia-related signature remained positively enriched in IC-EwS^>OPT^ subpopulation, compared to the rest of cells. Interestingly, several members of the HIF1α transcription factor network pathway including the anti-apoptotic hypoxia-induced gene *MCL1* were expressed significantly more in IC-EwS^>OPT^ than in IC-EwS^<OPT^ (Figure S5). This observation can indicate an EWSR1-FLI1-modulated interplay between hypoxia and apoptosis regulation in cells falling out of the optimal proliferation range of IC-EwS scores. In favor of this hypothesis, in the top 10 differentially expressed genes between IC-EwS^OPT^ and IC-EwS^<OPT^ cells there are several apoptosis-related genes, namely pro-apoptotic factor *BNIP3* (down-regulated in IC-EwS^OPT^) and Cytochrome C oxidase *COX6C* (up-regulated in IC-EwS^OPT^) (Figure S6).

We can conclude that the major sources of transcriptomic heterogeneity inside EwS PDXs are proliferation and the variation of activity of the EWSR-FLI1 transcription factor. We observe that there is an optimal range of activity of EWSR-FLI1 for cells to proliferate. We show that outside of this range, we can define cell subpopulations expressing hypoxia markers and genes involved in glucose pathways.

## Discussion

As most pediatric cancers, EwS is characterized by a paucity of genetic abnormalities. Accumulation of genetic alterations that frequently result from genetic instability, which are suspected to play major roles in the progression of adult cancers, is therefore not expected to be the major player in the progression of pediatric cancers.

Here, based on our recent findings that the variation of expression of *EWSR1-FLI1* constitutes a major source of heterogeneity in EwS, we used a variety of experimental systems to investigate at the single cell level the gene expression dynamics associated with changes in *EWSR1-FLI1* expression. To our knowledge this is the first report studying the dynamics of a transcriptome at the single cell level after induction of a known cancer driver gene.

We applied unsupervised independent component analysis (ICA), which first identified two components that are extremely specific to G1/S and G2/M cell cycle phases (IC-G2/M and IC-G1/S). These components are not EwS-specific and characterize a subset of cells in all the experimental systems included in the analysis. In EwS cells, IC-G2/M and IC-G1/S are clearly correlated with the expression level of *EWSR1-FLI1*. Furthermore an independent component exquisitely specific for EWSR1-FLI1 activity was identified which i) did not highlight any non-Ewing cells or tumors in single cell, tumor bulk or normal tissue datasets, ii) is strongly enriched by EF1 peaks associated with the presence of GGAA-mSats sequences in the vicinity of the TSS, iii) is strongly associated with EWSR1-FLI1-dependent super-enhancers regions. Based on filtering genes associated with this component, we further identified a set of 83 strong candidate genes for direct regulation by EWSR1-FLI1, a characteristic which was not reported previously for most of them. Previously proposed lists of EWSR1-FLI1 direct targets contain numerous cell cycle genes as a result of the correlation between *EWSR1-FLI1* induction and cell cycle gene expression. In contrast, this restricted list only contains *CCND1* as a gene directly involved in cell cycle strongly reinforcing the role of this gene as a major player in *EWSR1-FLI1-*induced activation of the cell cycle. Clear distinction between IC-EwS and IC-G2/M and IC-G1/S underlines the ability of the ICA approach to discriminate the cell cycle process, a usual confounding factor which hides the effects of other essential factors, and EWSR1-FLI1 direct downstream regulated genes. This is an alternative to the use of other methods that have been developed to “subtract” the signal related to the cell cycle signal from the data(Bacher and Kendziorski, 2016; Barron and Li, 2016), a step which would not be suitable in our study as the cell cycle and the proper oncogene transcriptional programs appear to be highly correlated.

The identification of the IC-EwS signature constitutes a considerable improvement as compared to the previously defined EWSR1-FLI1 signatures. When investigated with functional annotation tools (*Toppgene*, DAVID), IC-EwS only retrieves weak enrichment annotations as synapses, neurogenesis, or cell adhesion, in agreement with previous observations that EWSR1-FLI1 activates some neural and cell-cell adhesion processes (Franzetti et al., 2017; Hu-Lieskovan et al., 2005). Rather, this list contains genes involved in a variety of functions highlighting the pleiotropic effects of EWSR1-FLI1. Importantly, a number of these genes appears to be controlled by EWSR1-FLI1 binding to GGAA-mSats sequences that are highly polymorphic in the human population. The hypothesis of polymorphisms in these regions being involved in hereditary susceptibility, as shown recently for the *EGR2* locus (Grunewald et al., 2015), or in the inter-tumoral heterogeneity of EwS can now be more directly tested.

Exploration of 5 EwS tumors based on these independent components and on the most significant functional reference genesets they pointed at, enables to illuminate some aspects of intratumoral heterogeneity. One distinct group corresponds to actively proliferating cells. The number of cycling cells is variable, from 9% to 30%. Rather expectedly, these cells demonstrate increased scores for oxidative phosphorylation signatures.

We observe strong cell-to-cell variability of IC-EwS signature score, being an indirect measure of EWSR1-FLI1 activity. As expected, cells with a low IC-EwS score are not cycling, in agreement with the hypothesis of a significant expression of EWSR1-FLI1 being necessary for cell cycle entry and progression. More surprisingly, cells with the highest IC-EwS scores are not cycling either suggesting that proliferation of Ewing cells is induced by an intermediary, potentially optimal level of EWSR1-FLI1 expression (called IC-EwS^OPT^ in this study).

In addition to the cell cycle, EWSR1-FLI1 expression may induce metabolic heterogeneity. In all tumors, our analyses highlight a subgroup of IC-EwS^<OPT,>OPT^ cells that are characterized by hypoxia signal. We observe that the number of cells that may be assigned to a Warburg effect, *i*.*e*. an aerobic glycolysis, appears relatively low. It will be of strong interest to follow *in vivo* the relationship of these cell subpopulations with the microenvironment including blood vessels, fibroblasts and immune cells, and follow their evolution in response to therapy.

In conclusion, in this study we characterize the dynamic effect of EWSR1-FLI1 at single cell level. We can distinguish, in an unsupervised and unbiased manner, its oncogene-specific transcriptional program (IC-EwS) from a process strongly coupled to it, the induction of proliferation. The IC-EwS allowed us to describe tumoral heterogeneity in EwS’s PDXs highlighting three major populations: one corresponding to the optimal window for cell proliferation activation and two others characterized by lower or higher activity of IC-EwS and associated to hypoxia.

The methodology developed can be applied to other biological contexts, for example, in order to dissect different transcriptional programs of other known cancer drivers or tumor suppressors. The different components identified enable to characterize the transcriptional heterogeneity of a tumor cell population. Further studies will be needed to assess whether the composition of tumors in these different compartments influences the response to treatment and prognosis of tumors.

## Acknowledgements

We thank the personnel of the Unité de Génétique Somatique for their help to generate PDX models. This work is supported by ITMO Cancer SysBio program (MOSAIC project, Nb bio 2014-2: Modeling Single cell vAriability in Cancer progression). UK acknowledges the the Ministry of Education and Science of the Republic of Kazakhstan for the grant research project “Pan-cancer deconvolution of omics data using Independent Component Analysis” (IRN: AP05135430). This work was also supported by grants from the Institut Curie, the Inserm, the Ligue Nationale Contre le Cancer (Equipe labellisée), the ANR-10-EQPX-03 from the Agence Nationale de la Recherche, the European PROVABES (ERA-649 NET TRANSCAN JTC-2011), and ASSET (FP7-HEALTH-2010-259348) projects. This research was supported by FP7 grant “EURO EWING Consortium” No. 602856 and EU Horizon-2020 program (grant No 826121, iPC project) and the following associations: Courir pour Mathieu, Dans les pas du Géant, Les Bagouzamanon, Enfants et Santé, M la vie avec Lisa, Lulu et les petites bouilles de lune, les Amis de Claire, l’Etoile de Martin and the Société Française de lutte contre les Cancers et les leucémies de l’Enfant et de l’adolescent.

## Author Contributions

A.Z., and O.D. designed the study. M.M.A., O.M., S.G., N.G., A.Z. and O.D. wrote the manuscript. M.M.A, N.G., S.D., performed the single cell experiments and analyses. M.M.A, S.Z. and D.S. investigated the chromatin marks. N.G. and S.Z. performed *in vivo* experiments. NG, S.Z. and D.S. performed the xenograft and PDX experiments. O.M., A.Z., S.G., O.S, S.G., J.P.V., U.K., V.B. and E.B. performed the bioinformatics and the statistical analyses. V.R. M.B and S.B did all the sequencing experiments. F.T and I.J.L provide important materials or data used in this study. T.G.P established MSC lines.

## Methods

### Cell lines and animal experiments

All cells are grown at 37°C with 5% of CO_2_ with 100 UI/mL Penicillin and 100 µg/mL Streptomycin (Gibco). A673/TR/shEF (Carrillo et al., 2007) are cultured in DMEM (Gibco) 10% FBS (Eurobio), with 50 µg/mL Zeocin (Invitrogen), 2 µg/mL Blasticidin (Invitrogen) added *ex-tempo*. I2A cells were grown in RPMI (Gibco), 10% FBS (Eurobio) with 50 µg/mL hygromycin B (Life Technology), 300 µg/mL G418 (geneticin) and 50 ng/mL DOX (doxycyclin, Invitrogen) added *ex-tempo* when indicated. MSCs from bone marrow Ewing patients were isolated by density-gradient centrifugation using Ficoll technique and were cultured in alpha MEM (Gibco), MSC-qualified serum (Gibco), 1% L-glutamine (Gibco) and 1 ng/mL bFGF (Sigma), added *ex-tempo*. CLB-berlud are cultured in RPMI (Gibco) 10% FBS (Eurobio), 100U/mL.

*EWSR1-FLI1* specific small hairpin RNA was induced in A673/TR/shEF cells by adding DOX at 1 ug/mL. After 7 days, DOX was removed and cells were washed three times to allow silencing of the shRNA and induction of *EWSR1-FLI1*. Cells were harvested at seven different times points: 0 day (d7), 2 days (d9), 3 days (d10), 4 days (d11), 7 days (d14), 10 days (d17) and 15 days (d22) after DOX removal.

For A673/TR/shEF xenograft, 20 millions cells were resuspended in 200 µL of PBS and subcutaneously injected into severe combined immunodeficiency (SCID) mice. When tumor volume reached 1,000 mm3, DOX was added in the drinking water of a subset of mice (+ DOX group) for 7 days.

For I2A cells, DOX was removed and cells were washed three times to induce *SMARCB1* expression.

### Patient-derived xenograft

Ewing Patient Derived Xenografts (PDX) were established in the laboratory by subcutaneous implantation of tumor samples in SCID mice. Patients consented preoperatively to take part in the study which received agreement by the Institutional Review Board Protocol.

Eight PDX from EWSR1-FLI1-positive EwSs were profiled using either the Fluidigm™(PDX-83, PDX-84 and PDX-111) or the 10xGenomics (PDX-184, PDX-352, PDX-856, PDX-861 and PDX-1058) single cell technology. Four of them were derived from localized primary tumor, expressing an *EWSR1-FLI1* fusion type 1 transcript, either localized to the humerus and presenting a *CDKN2A* gene deletion (IC-pPDX-3 model: PDX-84 and PDX-184) or located in the sacrum and presenting a *STAG2* R614* gene mutation (IC-pPDX-8 model: PDX-83 and PDX-352). PDX-111 and PDX-861 were derived from an Ewing primary tumor presenting metastasis at diagnosis and expressing an *EWSR1-FLI1* fusion type 1 (IC-pPDX-52 model: PDX-861) or type X transcript (IC-pPDX-5 model: PDX-111). Finally, PDX-856 (IC-pPDX-80 model) and PDX-1058 (IC-pPDX-87 model, relapse from IC-pPDX-52 mode) were obtained from a relapse and expressed an *EWSR1-FLI1* fusion type 1 transcript. PDX-1058 presented a *CDKN2A* gene deletion and a *TP53* R175C* gene mutation.

### Tumor dissociation into single-cell suspension

A673/TR/shEF xenografts and Ewing PDX were dissected from mice and mechanically dissociated. The finely minced tissue was transferred to a digestion mix consisting of C0_2_ independent medium (Gibco) containing 1 mg/mL collagenase D (Roche), 2 mg/mL hyaluronidase (Sigma) and 25 µg/mL DNAse (Sigma), incubated for 45 min at 37 °C and gently resuspended every 10 min. Cell suspension was then filtered using 70 µm and 30 µm cell strainers (Miltenyi Biotec). For A673/TR/shEF xenograft experiments, the tumoral suspension was depleted of infiltrated murine cells using the mouse cell depletion kit from Milteny Biotec. Cells were then adjusted at 1×10^6^ cell/mL in HBSS containing 2 mM EDTA. Viability was measured using trypan blue exclusion.

### Western blot

All A673/TR/shEF *in vitro* and xenograft proteins were extracted with RIPA and anti-protease cocktail (Roche). Western blots were hybridized with rabbit monoclonal anti-FLI1 antibody (1:1000, ab133485, abcam) and mouse monoclonal anti-beta-actin (1:10,000, A-5316, Sigma Aldrich). Then, membrane was incubated with anti-mouse/rabbit IgG horseradish peroxidase coupled (1:3,000, Amersham Bioscience). Proteins were detected using chemiluminescence (Pierce).

### C1 single cell capture and mRNA-seq

Dissociated cells were captured and processed with the C1 Single-Cell Auto Prep System (Fluidigm™) following the manufacturer’s protocol. We started with a cell suspension at a concentration of 0.45 x 10^6^ cells/mL. After observation at the microscope, we identified the sites where live single cells were captured. Processing of cells occurred in the C1 instrument to perform steps of cell lysis, cDNA synthesis with reverse transcriptase, and PCR amplification for each cDNA library. Quality of the resulting cDNA was checked using the LabChip GX Touch HT (Perkin Elmer, Waltham, MA). The cDNA synthesis and PCR used reagents from the SMARTer Ultra Low RNA kit from Illumina sequencing (Clontech, Mountain View, CA). After harvest from the C1 device, each cDNA library was tagmented using the Nextera XT DNA Sample Preparation Kit (Illumina). After PCR, cDNA libraries were pooled. All libraries were sequenced with HiSeq2500 (Illumina) using 150 bp paired-end sequencing.

### 10x Genomics single cell capture and mRNA-seq

Single-cell RNA-seq was performed using the Single Cell 3’GEM Code Single-Cell instrument (10x Genomics, Pleasanton, CA, USA), according to the manufacturer’s protocol. Cellular suspension (5,300 cells) was loaded on a 10x Chromium instrument to generate 3,000 single-cell GEMs, using the Chromium™ Single Cell 3’ Library & Gel Bead Kit v2. The initial step consisted in performing an emulsion where individual cells were isolated into droplets together with gel beads coated with unique primers bearing 10x cell barcodes, UMI (Unique Molecular Identifiers) and poly(dT) sequences. GEM-RT was performed to generate barcoded full-length cDNA (53°C for 45 min, 85°C for 5 min, held at 4°C). After RT, GEMs were broken using the recovery agent and the single-strand cDNA was cleaned up with DynaBeads MyOne Silane Beads (Thermo Fisher Scientific).

Bulk cDNA was amplified (98 °C for 3 min; 12 cycles : 98 °C for 15 s, 67 °C for 20 s, 72 °C for 1 min; 72 °C for 1 min; held at 4 °C) and then cleaned up with the SPRIselect Reagent Kit (Beckman Coulter). A qualitative analysis on the amplified cDNA was performed using Agilent Bioanalyzer high sensitivity chip.

Libraries were then constructed in 4 main steps: 1) fragmentation, end repair and A-tailing, 2) adaptor ligation, 3) post ligation cleanup with SPRIselect and 4) sample index PCR, quantified by Qubit fluorometric assay (Invitrogen) with dsDNA HS (High Sensitivity) Assay Kit and qualified using LabChip (LabChip® GX Touch™ PerkinElmer).

Indexed libraries were equimolarly pooled and sequenced on an Illumina HiSeq 2500 in rapid run mode, using paired-end (PE) 26/98 according to 10x recommendations. Using a full rapid flow cell, a coverage around 100M reads per sample were obtained corresponding to 100,000 reads/cell.

In order to remove profiles corresponding to dead or stressed cells from the analysis of 10x data, mitochondrial percentage score was computed for each cell as the percentage of UMIs captured by the genes from the previously described gene set (Ilicic et al., 2016). In the histograms of this score, a bimodal distribution was observed; therefore, all cells from the higher mode were removed from the analysis. After additional quality checks such as removal of cells with too small total number of UMIs (<5,000 UMIs per cell, compared to the median 15,000 number of UMIs per cell) or too high (>40,000 UMIs), the number of selected cells in each PDX is indicated in the Results section.

### Alignment, counting and sample normalization of reads

Reads obtained from sequencing of cells were aligned on the human genome (v. hg19) using TopHat (version 2.0.6) (Trapnell et al., 2009). Reads mapping more than once (parameter -x 1) or having edit distances of more than 3 (-N 3) were discarded.

Counting of reads on annotated genes from the GRCh37 gene build was done using htseq-count (v. HTSeq-0.5.3p9) (Anders et al., 2015) with the following parameters: reads with a quality score less than 10 (-a 10) were discarded and reads partially overlapping with the annotated gene transcript were included in the counts unless they overlapped with another read. In all experiments analysed the STRANDED=no option was used.

Sample-to-sample normalization was performed by rescaling using DESeq size factors (Love et al., 2014). For all data analyses the number of reads was log10(x+1) transformed.

In case of 10x Genomics data, the programs “cellranger mkfastq” and “cellranger count” from the Cell Ranger software suite (v. 1.3.1) provided by 10X Genomics were used for demultiplexing and counting the reads on the reference genome GRCh38. Sample-to-sample normalization was performed using the total number of reads in the log scale. For all data analyses the number of reads was log_10_(x+1) transformed. More specifically if X is the count matrix the R code to obtain the normalised matrix X.tpm is the following: median.umi <- median(colSums(X)); X.tpm <- log(t(t(X)/colSums(X))*median.umi+1). For each cell, reads from the k = 5 most similar cells were pooled together to define the new cell measurement, in order to reduce the effect of drop-outs. For pooling, kNN graph was computed on log10(x+1) transformed data after filtering non-variant genes (variance smaller than 0.01) and reducing the dimension of the data by projecting it into 20-dimensional subspace spanned by the standard PCA components.

### Exploratory analysis of scRNA-seq data

ICA was applied as previously described (Biton et al., 2014), using stabilization, with an additional procedure for determining the optimal number of independent components (Kairov et al., 2017). In the ICA decomposition X = AS, X is the gene expression (sample *vs*. gene) matrix, A is the (sample *vs*. component) matrix describing the loadings of the independent components, and S is the (component *vs*. gene matrix) describing the weights (projections) of the genes in the components. We used a modified MATLAB implementations of fastICA (Hyvarinen, 1999) and icasso (Himberg et al., 2004) as a part of the BIODICA software available at https://github.com/LabBandSB/BIODICA/, which contains an algorithm for estimating the optimal number of components to compute. Icasso applies FastICA algorithm for finding *m* independent components n = 100 times, and then uses hierarchical clustering to estimate compactness of clusters of the components computed in all runs. The resulting independent components represent medoids of the *m* clusters and are ranked by the reproducibility (cluster compactness) in *n* runs. The orientation of the components was chosen such that the longest tail of the gene projection distributions would correspond to the positive values.

In order to explore the relation between IC metasamples and the binary vectors representing various subsets of cells, we used the mutual correlation metrics in which each IC was characterized by a vector of correlations with all other IC metasamples and the binary vectors (Table S2), normalized to the unity L1-norm. Subsequently, a standard PCA analysis was applied (Figure 2A).

t-SNE analysis(Van der Maaten, 2008) was done using R with setting the initial dimension parameter to 100 and the perplexity parameter set to 80.

SPRING visualization was produced through computing the kNN graph (*k* = 5) by applying a standard for SPRING approach (Weinreb et al., 2018) consisting in: 1) filtering genes with the coefficient of variance smaller than 0.05 and the average expression smaller than 0.01, computed for pooled read counts (this filter left from 8 to 9 thousands of genes in our datasets); 2) normalizing the measurements on the library size and 3-reducing the dimension of the dataset to 20 by the standard PCA algorithm.

### Functional enrichment analysis and gene set scoring

For interpreting the biological meaning of the sets of top-contributing to each of the ICs genes, we applied toppgene functional analysis tool (Chen et al., 2009), limited to reference gene sets not larger than 500 genes (in order to focus on more specific functional categories). The toppgene analysis was automated through BIODICA graphical user interface available from https://github.com/LabBandSB/BIODICA/ and recapitulated in the form of an interactive online table http://bioinfo-out.curie.fr/projects/sitcon/mosaic/toppgene_analysis/. The table is organized in two columns reporting the first most enriched functional gene sets for positive and negative part of each IC metagene, in each reference categories (the enriched function is mentioned in the table only if the the Bonferroni-corrected *p* < 0.05 and the number of genes from the function found in the top-contributing list is not smaller than 8). Also the sets of top-contributing genes smaller than 10 were not considered for the enrichment analysis. Each hyperlink in the form of “ICX+/-” leads to a saved detailed enrichment analysis as it was produced by toppgene. Each hyperlink in the form “X genes” leads to the tested list of top-contributing genes.

The table was further used to select a set of reference signatures for the analysis of the tumor data. Only signatures from GO and Pathway categories enriched with the Bonferroni-corrected *p* < 10^−10^ were selected for further analysis. On top of this, we added the standard HALLMARK set of transcriptomic signatures from MolSigDB.

For associating IC-EwS score computed for tumor cells with the reference signatures, we applied the standard pre-ranked GSEA analysis to the *t*-test scores computed between the 10th and 90th percentiles of the IC-EwS score. Classical scoring scheme was used and 1,000 permutations estimating the empirical *p*-value.

Gene set scores for gene sets were computed in all analyses as average gene expression of the genes composing the signatures, after removal of genes characterized by small variance (in all analyses, 2000 most variable genes were kept for computing the scores).

### Non-regulated control gene sets

We selected “Non-regulated genes” for which at least 100 reads were detected at d7 and/or d22 and which showed no significant differential expression between d7 and d22 in A673/TR/shEF bulk expression dataset (0.5 < FC < 2, *p* > 0.01) (n = 2,117). Then, genes were ranked from the lowest to highest fold change. For our analysis, we used the top 100 non-regulated genes for the Figure 2C and the top 1,000 non-regulated genes for the Figure 3 as a negative control.

### Chromatin-immunoprecipitation and sequencing

DNA-protein cross-linking was performed in the presence of 1% of paraformaldehyde on 12×10^6^ cells for each condition during 10 minutes. Cell lysis, chromatin shearing, immunoprecipitation and DNA purification was performed with reagents from iDeal ChIP-seq kit for Transcription Factors (Diagenode, ref: C01010054). Twenty cycles of sonication (30s high, 30s off) using TPX tube (Diagenode, ref: 50001) and the Bioruptor (Diagenode) were achieved for chromatin shearing. We took 2 µg of FLI1 rabbit polyclonal antibody (abcam, ab15289-ChIP grade) to perform immunoprecipitation of EWSR1-FLI1 transcription factor and 1 µg of H3K27ac antibody (abcam, ab4729) for histone mark immunoprecipitation. IgG and CTCF ChIP was included as negative and positive control. To check quality of each ChIP reactions, quantitative PCR was realized prior to sequencing on 1/5 of purified DNA. Tested regions correspond to following primers:1-*CCND1* (5’GGTGGGAGGTCTTTTTGTTTC3’/5’CACGCAATCCCAGATCAAAAC3’); 2-*CDKN1A* (5’ACTGACTCATCACTACTCCCTC3’/5’GTGTGCTATTCCCGCCAG3’); 3-*CCND1* (5’CACAGTGTGGGTATTTCCATCAAGCA 3’/5’GGTGTGTAGGAAAAACAGCTCTCTGGA3’);4-*Sec14L2* (5’GCCCCCGCTGATGCACTTCC3’/5’AAGTGCGCCAGCAGAGCCAG3’). ChIP and input were sequenced with HiSeq2500 (Illumina) using 100 bp single-end sequencing.

### ChIP-seq peak detection and annotation

ChIP-seq reads were aligned to the human genome (hg19 version) with Bowtie2(Langmead and Salzberg, 2012). Peaks were called with MACS2 (Zhang et al., 2008), with option narrow for FLI1 antibody and broad for H3K27ac histone mark. To normalize, we took the input dataset from the same cell line. EWSR1-FLI1 specific peaks were defined as peaks varying upon EWSR1-FLI1 expression (*p* < 0.005). To obtain the *p*-values for each of the peaks we tested the statistical correlation (lm function in R) between the vectors formed by the EWSR1-FLI1 peak intensities at d7, d9, d10, d11, d14, d17 and the vector c (0, 2, 3, 4, 7, 10). That last vector consists in the number of days of EWSR1-FLI1 re-expression for each of these time points. For each gene, we reported the closest EF-peaks to TSS. We performed a Wilcoxon test to compare the distribution of distances for genes of each IC with the control gene set (Figure 3A). We used FIMO tool (Grant et al., 2011) to scan EF-peaks with ETS motif (JASPAR ID: MA0475.1, *p* < 0.1) and GGAA-mSats (JASPAR ID: MA0149.1, *p* < 0.0005). If several motifs were found, we kept only the best motif. ROSE was used to predict Super-Enhancers from H3K27ac marks (Loven et al., 2013){Whyte, 2013 #77. We applied Fisher’s exact test to evaluate the enrichment of EF-peaks in Super-Enhancer (Figure 3D). The Super-Enhancers were associated to the closest expressed gene (Figure 3E-F).

## Supplementary Information

**Table S1.**
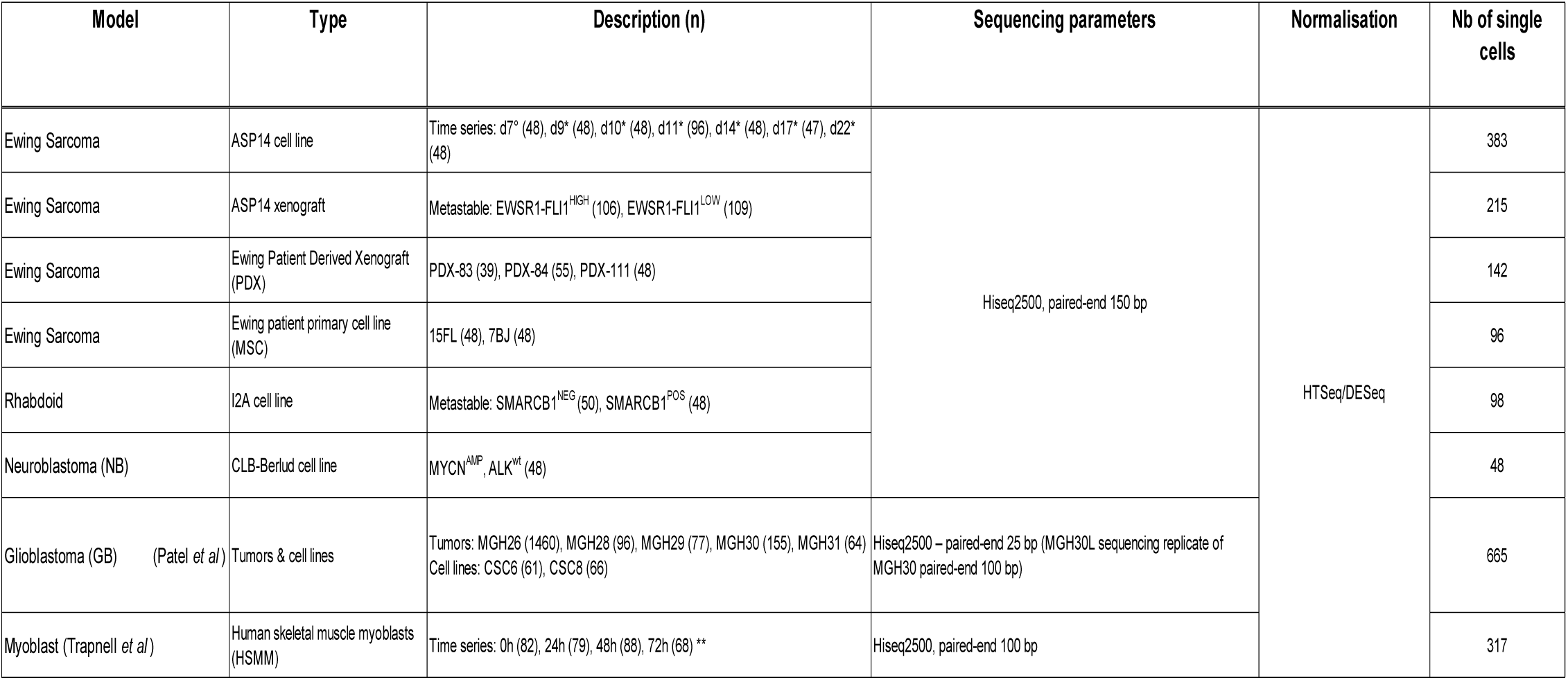
Summary of scRNA-seq datasets used in this study. (A673/TR/shEF, PDX, MSC, I2A, CLB-Berlud, glioblastoma and myoblasts). The model, the type of biological materials, the experiment description, the sequencing parameters, the normalization applied and the number of single cells sequenced are described for each dataset.

**Table S2.**
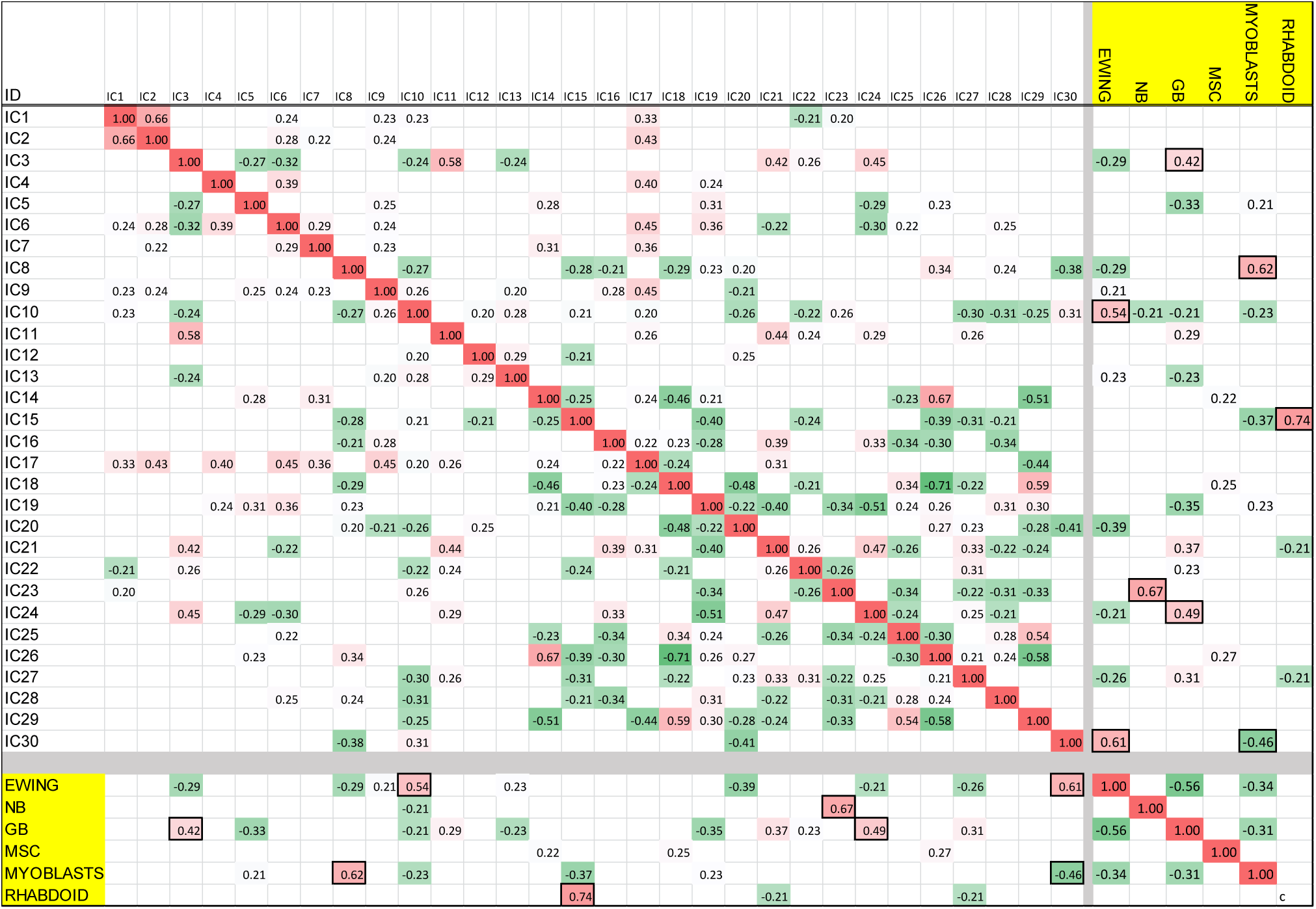
Correlations of the 30 components to the binary vector describing different cell subset.

**Figure S1.**
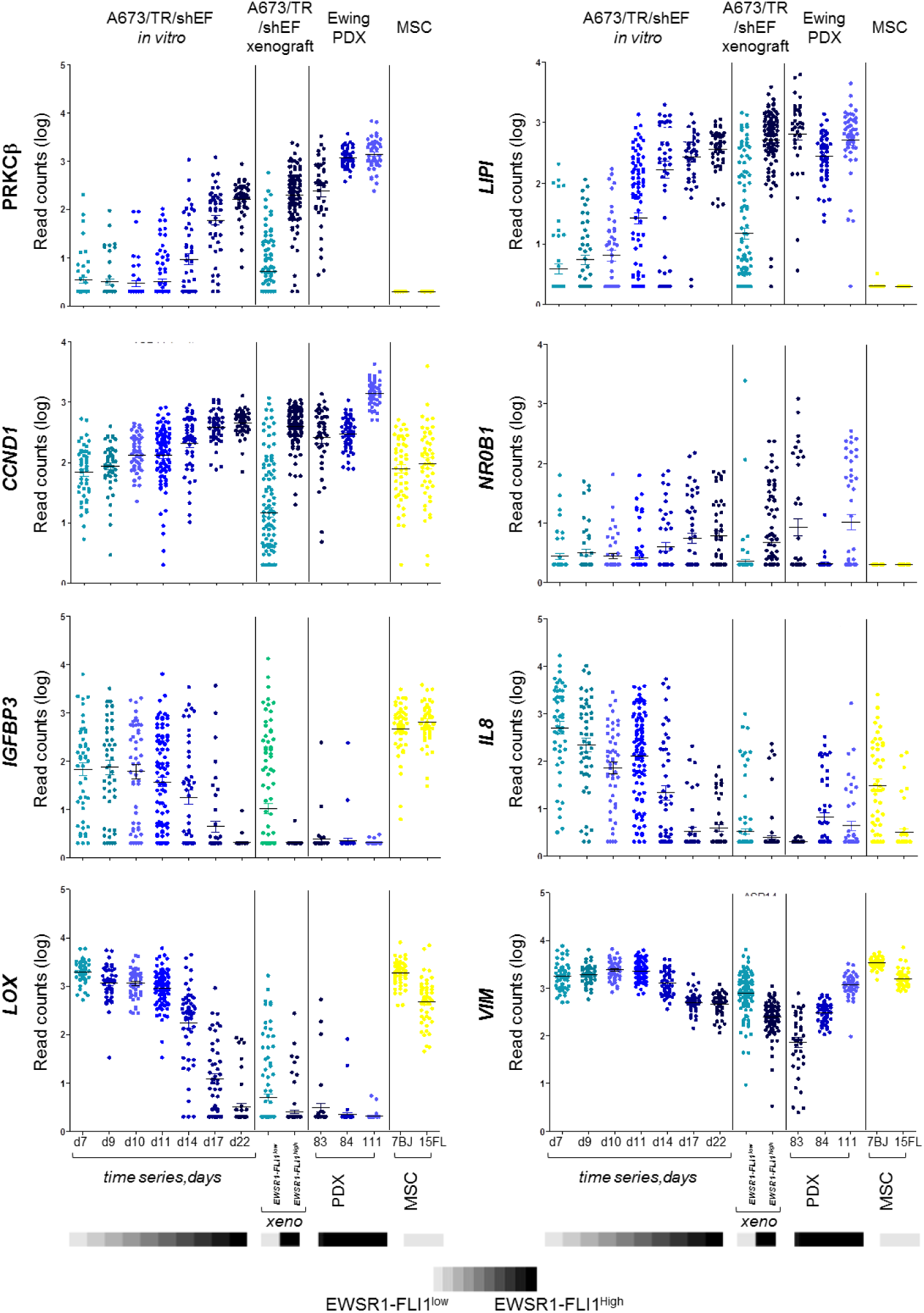
Expression of selected known regulated genes in EwS single cell dataset. Eight genes known to be modulated by EWSR1-FLI1 are plotted for A673/TR/shEF *in vitro*, A673/TR/shEF xenograft, PDXs and MSCs: 4 known up-regulated genes (*PRKCβ, LIPI, CCND1* and *NR0B1*) and 4 known down-regulated genes (*IGFBP3, IL8, LOX* and *VIM*). The grey scale on the bottom illustrates the EWSR1-FLI1 level of expression.

**Figure S2.**
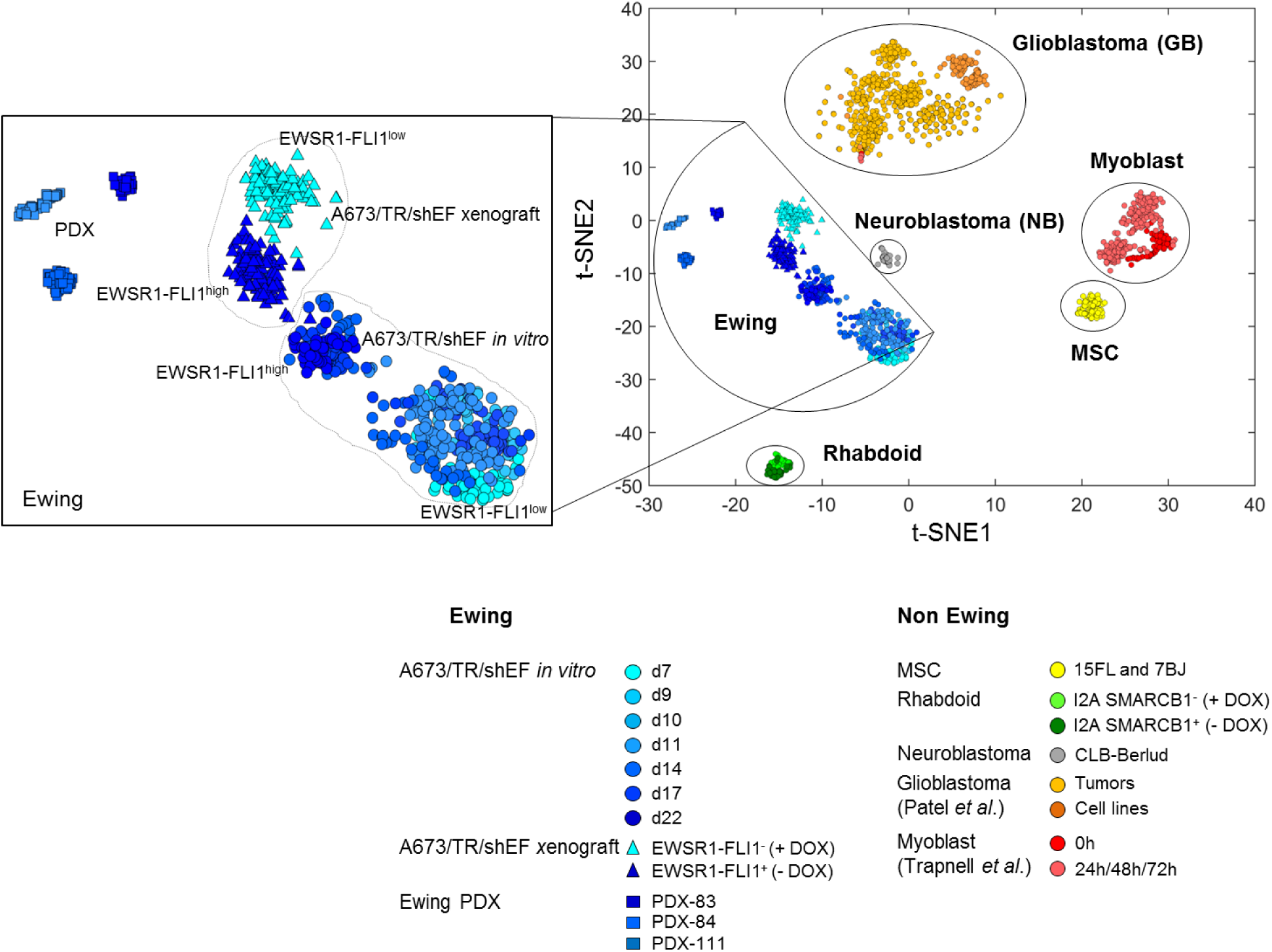
Exploratory analysis of several scRNA-seq datasets. t-SNE plot of the merged single cell data (1,964 profiles, 8 datasets jointly normalized, Table S1).

**Figure S3.**
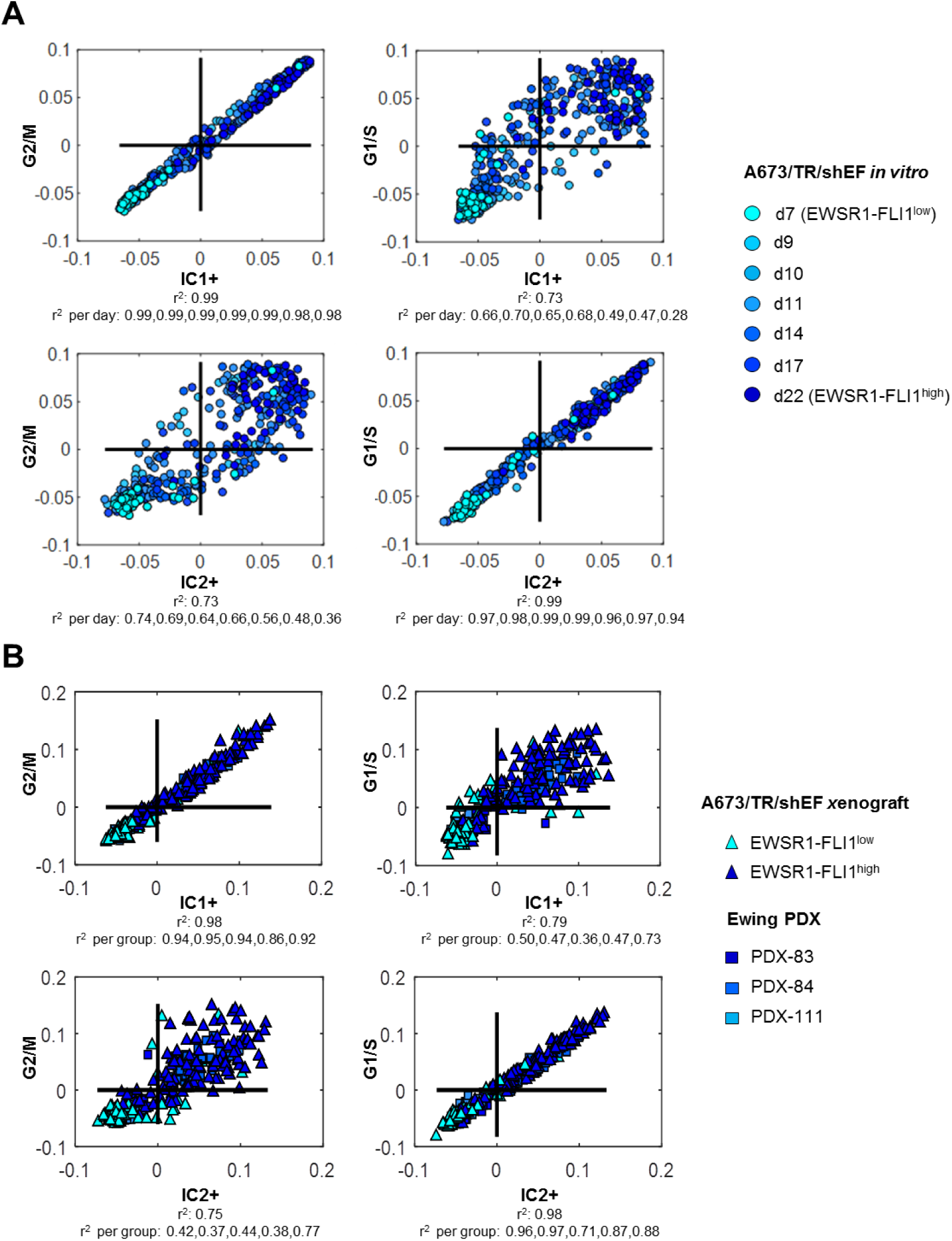
Comparison of cell cycle-associated independent components with cell cycle phase-specific transcriptomic signatures (G1/S and G2/M). Two scores characterizing progression of a cell through G1/S and G2/M phases of cell cycle computed using transcriptomic signatures obtained from meta-analysis of public datasets{Giotti, 2017 #57} are compared against IC1+ and IC2+ scores. Global correlation and correlation per time point or sample are indicated under each plot **A.** Pairwise scatterplots for the G1/S and G2/M scores *vs*. IC1+ and IC2+ scores allows matching IC1+ to G2/M score and IC2+ to G1/S score for A673/TR/shEF *in vitro* cells. **B.** The same comparison made in A673/TR/shEF xenograft cells and PDXs. The Pearson correlation coefficient is shown below the plots. The r^2^ per time point or per group are indicated underneath each correlation graph.

**Figure S4.**
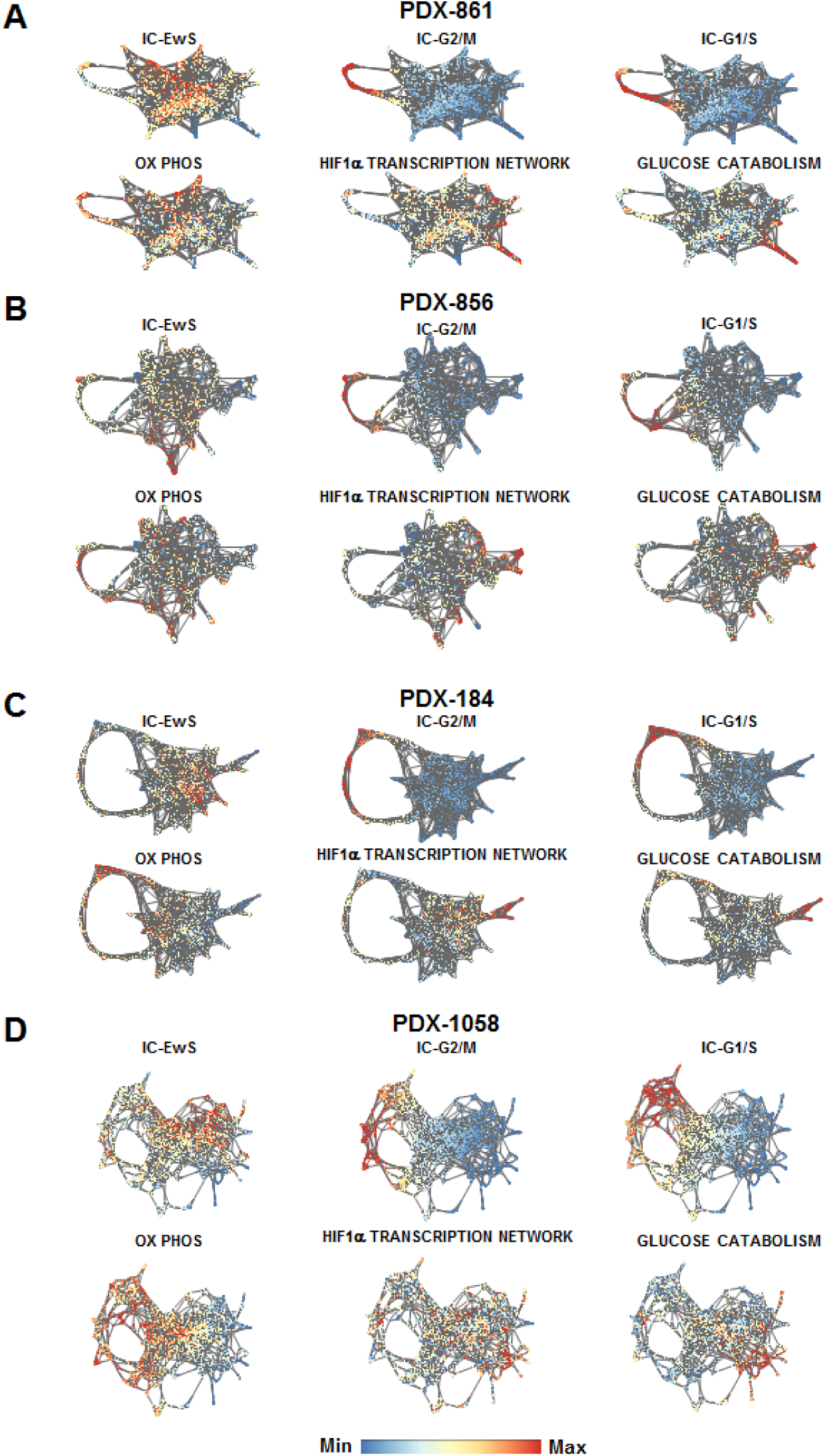
Analysis of tumors heterogeneity using force-directed layout of the similarity graph (SPRING software). **A-D**. SPRING representation of the kNN graphs (k = 5) for the four EwS PDXs: PDX-861, PDX-856, PDX-184 and PDX-1058 datasets respectively. The scores are either gene set (top contributing genes) scores of the ICs computed for model systems (IC-EwS recapitulating the transcriptional program of EWSR1-FLI1, IC-G2/M and IC-G1/S recapitulating the transcriptional programs of the cell cycle) or the scores of the reference gene sets recapitulating a biological function contributing to the intratumoral heterogeneity.

**Figure S5.**
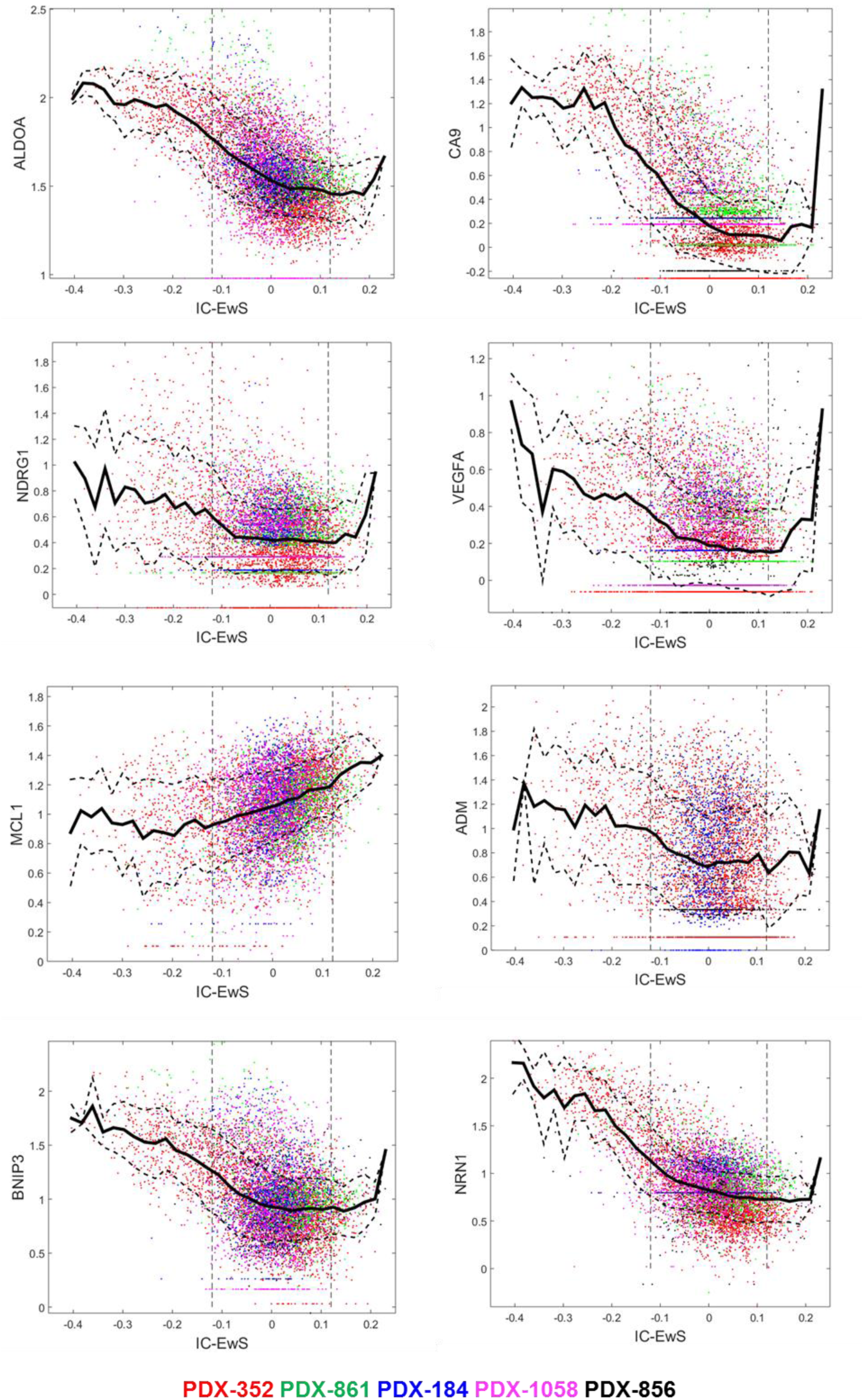
Expression of HIF1α transcriptional factor network of genes. **A-D**. Plot showing connection between IC-EwS and genes of HIF1α transcription factor network score for the five EwS PDXs. The expression of genes was rescaled in each PDX in order to have the same median expression. Solid line shows local average of the gene expression, and the dashed lines indicate one standard deviation interval. Only the data for the genes among the top 10k most variant genes in each PDX is shown.

**Figure S6.**
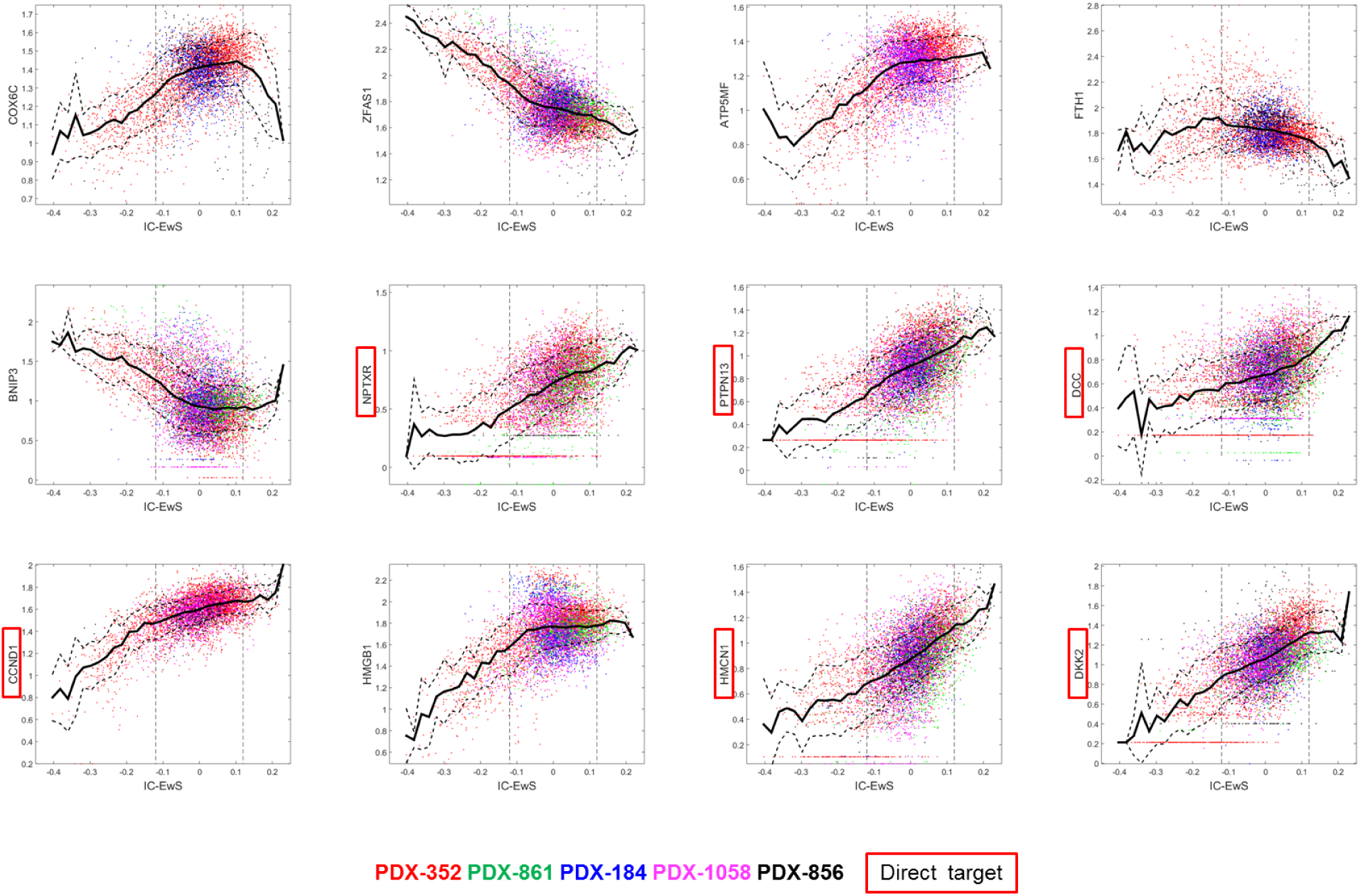
Expression of the top differentially expressed genes between IC-EWS^OPT^ and IC-EWS^<OPT,>OPT^. Plot showing connection between IC-EwS and several top genes differentially expressed between the optimal for proliferation region IC-EWS^OPT^ and other cell populations, for the five EwS PDXs. Solid line shows local average of the gene expression, and the dashed lines indicate one standard deviation interval. Only the data for the genes among the top 10k most variant genes in each PDX is shown.

